# Replicative Instability Drives Cancer Progression

**DOI:** 10.1101/2022.05.06.490945

**Authors:** Benjamin B. Morris, Jason P. Smith, Qi Zhang, Zhijie Jiang, Oliver A. Hampton, Michelle L. Churchman, Susanne M. Arnold, Dwight H. Owen, Jhanelle E. Gray, Patrick M. Dillon, Hatem H. Soliman, Daniel G. Stover, Howard Colman, Arnab Chakravarti, Kenneth H. Shain, Ariosto S. Silva, John L. Villano, Michael A. Vogelbaum, Virginia F. Borges, Wallace L. Akerley, Ryan D. Gentzler, Richard D. Hall, Cindy B. Matsen, C. M. Ulrich, Andrew R. Post, David A. Nix, Eric A. Singer, James M. Larner, P. Todd Stukenberg, David R. Jones, Marty W. Mayo

**Affiliations:** Department of Biochemistry and Molecular Genetics, University of Virginia, Charlottesville, Virginia, USA; Department of Pathology, University of Virginia, Charlottesville, Virginia, USA; Center for Public Health Genomics, University of Virginia, Charlottesville, VA, USA; M2Gen, Tampa, Florida, USA; Division of Medical Oncology, Department of Internal Medicine, Markey Cancer Center, Lexington, Kentucky, USA; Division of Medical Oncology, Department of Internal Medicine, The Ohio State University Comprehensive Cancer Center, Columbus, Ohio, USA; Department of Thoracic Oncology, H. Lee Moffitt Cancer Center and Research Institute, Tampa, Florida, USA; Division of Hematology/Oncology, Department of Internal Medicine, University of Virginia Comprehensive Cancer Center, Charlottesville, Virginia; Department of Breast Oncology, H. Lee Moffitt Cancer Center and Research Institute, Tampa, Florida, USA; Huntsman Cancer Institute and Department of Neurosurgery, University of Utah, Salt Lake City, Utah, USA; Department of Radiation Oncology, The Ohio State University Comprehensive Cancer Center, Columbus, Ohio, USA; Department of Malignant Hematology, H. Lee Moffitt Cancer Center and Research Institute, Tampa, Florida, USA; Department of Cancer Physiology, H. Lee Moffitt Cancer Center and Research Institute, Tampa, Florida, USA; Department of NeuroOncology, H. Lee Moffitt Cancer Center, Tampa, Florida, USA; Division of Medical Oncology, University of Colorado Comprehensive Cancer Center, Aurora, Colorado, USA; Department of Medical Oncology, Department of Internal Medicine, Huntsman Cancer Institute, Salt Lake City, Utah, USA; Department of Surgery, Huntsman Cancer Institute, University of Utah, Salt Lake City, Utah, USA; Huntsman Cancer Institute and Department of Population Health Sciences, University of Utah, Salt Lake City, Utah, USA; Department of Biomedical Informatics and Huntsman Cancer Institute, University of Utah, Salt Lake City, Utah, USA; Department of Oncological Sciences, Huntsman Cancer Institute, Salt Lake City, Utah, USA; Section of Urologic Oncology, Rutgers Cancer Institute of New Jersey, New Brunswick, New Jersey, USA; Department of Radiation Oncology, University of Virginia Comprehensive Cancer Center, Charlottesville, Virginia, USA; Department of Thoracic Surgery, Memorial Sloan-Kettering Cancer Center, New York, New York, USA

**Keywords:** Replicative instability (RIN), cancer progression, metastasis, MYBL2, single-strand break repair, translesion synthesis

## Abstract

In the past decade, defective DNA repair has been increasingly linked with cancer progression. Human tumors with markers of defective DNA repair and increased replication stress have been shown to exhibit genomic instability and poor survival rates across tumor types. Here we utilize-omics data from two independent consortia to identify the genetic underpinnings of replication stress, therapy resistance, and primary carcinoma to brain metastasis in BRCA wildtype tumors. In doing so, we have defined a new pan-cancer class of tumors characterized by replicative instability (RIN). RIN is defined by genomic evolution secondary to replicative challenge. Our data supports a model whereby defective single-strand break repair, translesion synthesis, and non-homologous end joining effectors drive RIN. Collectively, we find that RIN accelerates cancer progression by driving copy number alterations and transcriptional program rewiring that promote tumor evolution.

**Statement of Significance:** Defining the genetic basis of genomic instability with wildtype BRCA repair effectors is a significant unmet need in cancer research. Here we identify and characterize a pan-cancer cohort of tumors driven by replicative instability (RIN). We find that RIN drives therapy resistance and distant metastases across multiple tumor types.

## Introduction

Large scale sequencing efforts have enabled the discovery of genetic events that drive cancer development (1–5). Analysis of sequencing data has helped establish molecular classifiers through which tumors can be grouped according to mutations, copy number changes, or fusions. Careful study of genetic drivers has greatly expanded our understanding of how cancers develop. Despite this knowledge, primary tumors with similar genetic backgrounds often have highly heterogeneous outcomes, suggesting that there are additional factors that influence patient outcomes beyond initial oncogenic events. This is especially true when considering key cancer progression events such as therapy resistance and metastasis.

Decades of research has revealed processes dysregulated by cancers, summarized as hallmarks of cancer (6–7). Of these hallmarks, genomic instability has been linked to progressive disease across tumor types (8). Familial genetic studies and cancer genome analyses have revealed that genomic instability can develop following inactivation of DNA repair genes, such as BRCA1, BRCA2, and BRCA-related genes (9). As a result, tumors rely on error-prone DNA repair pathways and accumulate mutations and chromosomal alterations (9). Clinically, tumors with defective BRCA genes can be targeted using PARP inhibitors and platinum chemotherapies (10). However, it is recognized that genomic instability is observed in tumors that lack BRCA inactivation (11). Importantly, these tumors respond poorly to PARP inhibition, chemotherapy, and irradiation (11). Defining the genetic underpinnings of tumors with genomic instability and wildtype repair effectors is a significant unmet need in cancer research.

Previous work from our group revealed that elevated expression of the transcription factor MYB proto-oncogene like 2 (MYBL2) identified lung adenocarcinomas with genomic instability and wildtype BRCA (12). Our initial studies revealed that this *MYBL2* High phenotype was associated with a unique set of cancer genetics and identified patients at risk for poor outcomes. In this manuscript, we sought to identify a pan-cancer mechanism that underpins genomic instability and cancer progression in tumors with wildtype BRCA. In this study, we provide evidence that elevated *MYBL2* expression is a robust marker of poor patient outcomes across tumor types and genotypes. Importantly, this *MYBL2* High cohort is defined by genomic instability and inefficient homologous recombination despite containing wildtype BRCA. Analysis of the DNA repair landscape revealed that the underlying genetic basis of *MYBL2* High disease are heterozygous losses of single-strand break repair, translesion synthesis, and/or non-homologous end-joining effectors. These genetic lesions cause *MYBL2* High tumors to experience significant replication stress. Functional clustering of replication stress sensitive sites revealed that elevated replication stress promotes copy number alterations that rewire transcriptional programs and impact hallmarks of cancer master regulators. Clinically, this phenotype identifies patients at risk for poor outcomes when treated with chemotherapy and irradiation. Furthermore, our results demonstrate that *MYBL2* expression stratifies patient risk for distant metastases, especially to the brain. Our data defines a new pan-cancer class of tumors driven by replicative instability (RIN), unifying seemingly disparate cancers. Moreover, these results define a new mechanism through which RIN accelerates cancer progression by impacting several hallmarks of cancer.

## Results

### Pan-cancer analysis identifies *MYBL2* expression as a robust marker of poor patient outcomes across tumor types and genotypes

To test if *MYBL2* expression identified patients with poor outcomes and progressive disease across tumor types, we analyzed 32 studies curated by The Cancer Genome Atlas (TCGA) and other groups (13). For each study, samples were stratified based on *MYBL2* mRNA expression using a quartile approach (**Figure 1A**). To be included in further analyses, *MYBL2* expression had to identify patients with significantly inferior overall survival (OS) and progression free survival (PFS) outcomes. Kaplan-Meier analyses confirmed that *MYBL2* expression was a robust marker of poor patient outcomes in multiple tumor types, including lung adenocarcinoma (LUAD), isocitrate dehydrogenase (IDH)-mutant lower grade glioma (IDH^MUT^ LGG), pancreatic adenocarcinoma (PAAD), uterine corpus endometrial carcinoma (UCEC), and sarcoma (SARC) (**Figure 1B, Supplementary Table 1**). Across these tumor types, patients with *MYBL2* High disease had significantly worse OS, disease-specific survival (DSS), and PFS outcomes compared to patients with *MYBL2* Low tumors.

**Figure 1:**
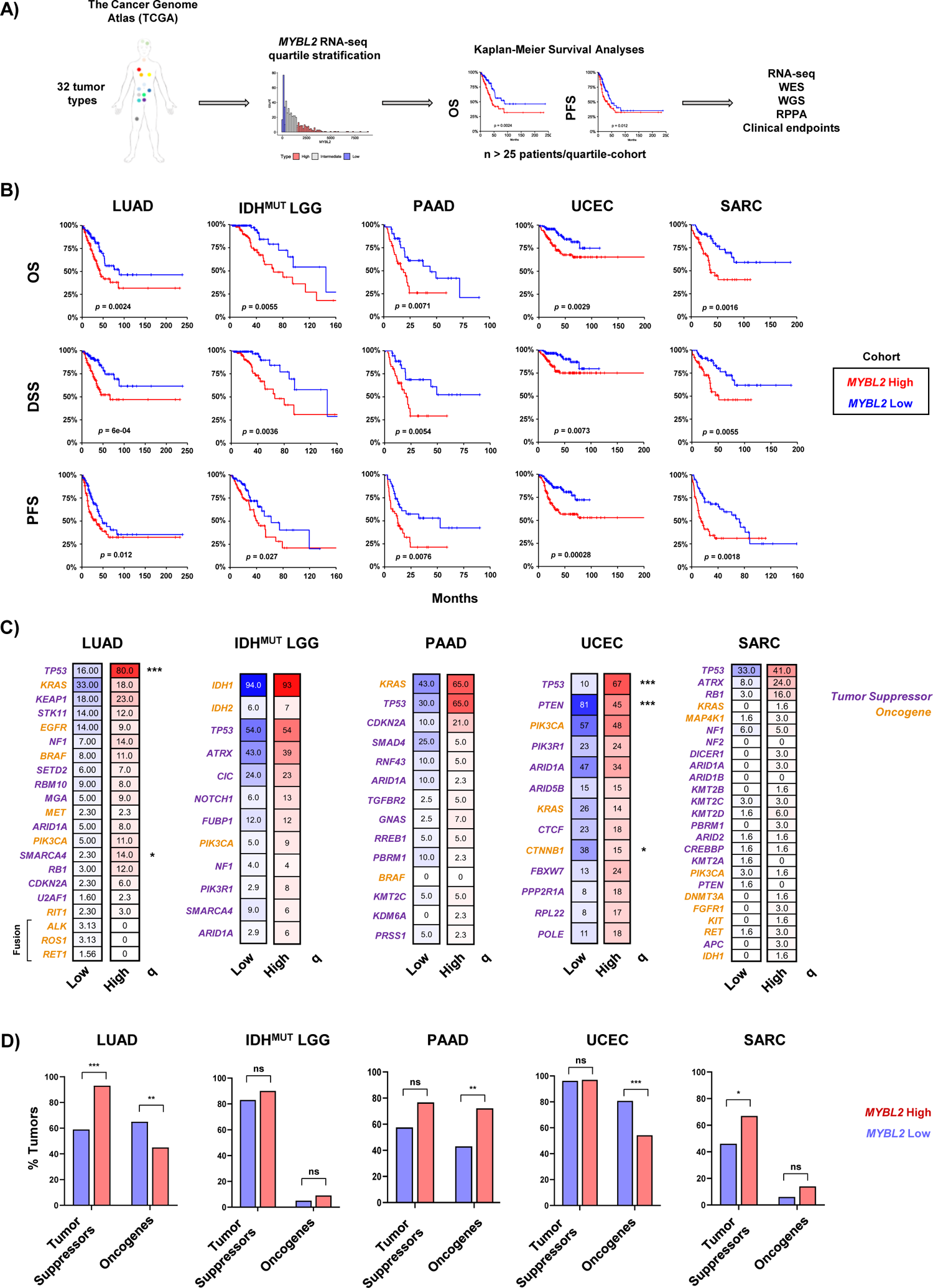
Elevated *MYBL2* mRNA expression identifies patients with poor outcomes across multiple tumor types and genotypes. A) Pan-cancer analysis overview. **B)** Kaplan-Meier analyses demonstrate that *MYBL2* expression is robustly prognostic across multiple tumor types for OS, DSS, and PFS outcomes. Log-rank test *p*-values are displayed. C) *MYBL2* High tumors develop across common cancer genetic driver backgrounds. Percentages reflect the percent of tumors with gene specific alterations. Statistical significance mapping represents Benjamini-Hochberg corrected *q* values, *q* < 0.05 *, *q* < 0.01 **, *q* < 0.001 ***. **D)** Individual tumors show different patterns of tumor suppressor inactivation and oncogene activation with respect to *MYBL2* High and *MYBL2* Low disease. IDH^MUT^ LGG tumor suppressor and oncogene status were mapped excluding founding IDH mutations. One-sided Fisher’s exact test, *p* < 0.05 *, *p* < 0.01 **, *p* < 0.001 ***. LUAD: lung adenocarcinoma. IDH^MUT^ LGG: IDH-mutant lower grade glioma. PAAD: pancreatic adenocarcinoma. UCEC: uterine corpus endometrial carcinoma. SARC: sarcoma.

Next, *MYBL2* High and Low tumors were profiled for tumor specific genetic driver events as defined previously (**Figure 1C**) (1–5). Surprisingly, this analysis demonstrated that *MYBL2* High tumors develop across common cancer genotypes, with few statistically significant enrichments for individual driver alterations. Notable exceptions include enrichments for TP53 and SMARCA4 mutations in LUAD and TP53 mutations in UCEC. *MYBL2* High UCEC was also inversely correlated with PTEN and CTNNB1 mutations. Given the lack of enrichment, driver genes were binned into broad tumor suppressor and oncogene categories to test for general enrichment patterns (**Figure 1D**). This analysis also failed to identify a clear pattern, indicating that there are additional steps beyond known driver mutations that are required to generate *MYBL2* High tumors.

### *MYBL2* High tumors are characterized by genomic instability and inefficient homologous recombination despite containing wildtype BRCA

To characterize similarities of *MYBL2* High disease, we analyzed DNA damage metrics provided by the TCGA PanCancer working group (13, 14). Using these data, we found *MYBL2* High tumors universally had significantly elevated mutation burden as well as greater fractions of the genome altered (FGA) (**Figure 2A**). All *MYBL2* High tumor cohorts exhibited significantly greater levels of microsatellite instability (MSI) (**Figure 2B**). It should be noted that only a small number of samples across tumor types reach the threshold required to be deemed ‘MSI-High’ (MSISensor score ≥ 10), most of which are UCECs (15). Regardless, separating tumors based on *MYBL2* mRNA expression consistently identified tumors with varying degrees of elevated MSI. Taken together, these data demonstrate that genomic instability is a hallmark of *MYBL2* High disease.

**Figure 2:**
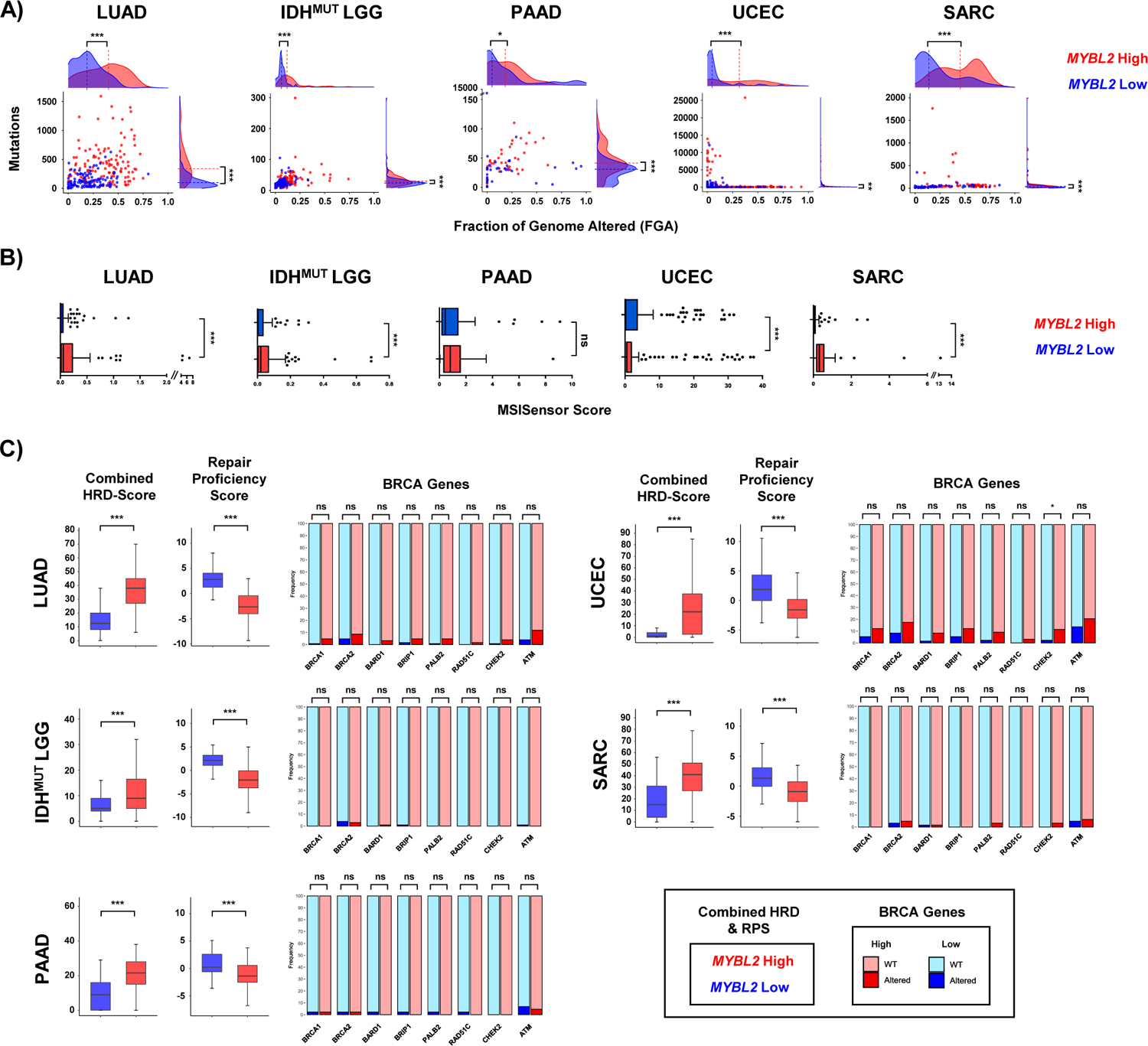
*MYBL2* High tumors exhibit genomic instability despite containing wildtype BRCA genes. **A)** *MYBL2* High tumors have significantly greater somatic mutation and fraction of the genome (FGA) altered. **B)** *MYBL2* High tumors have elevated microsatellite instability scores. **C)** *MYBL2* High tumors exhibit inefficient homologous recombination despite containing wildtype BRCA genes. **A), B), C)** Statistical significance was assessed using Wilcoxon signed rank tests (). *p* < 0.05 *, *p* < 0.01 **, *p* < 0.001 ***. **C)** Enrichments for inactivating alterations in BRCA genes were tested using one-sided Fisher’s exact tests. Significance is mapped using Benjamini-Hochberg corrected *q* values. *q* < 0.05 *, *q* < 0.01 **, *q* < 0.001 ***; ns, not significant.

Studies have shown that a common cause of genomic instability is a loss of homologous recombination (HR) repair (9). To analyze the status of HR repair, we analyzed combined homologous recombination deficiency (combined HRD) scores and repair proficiency scores (RPS) (14, 16). Combined HRD scores are derived from the presence of genomic scars as they reflect the sum of chromosomal alterations impacting telomeric regions, loss of heterozygosity events, and large-scale transitions (14). Tumors with high combined HRD scores exhibit elevated genomic instability. The repair proficiency score is an RNA-based metric that captures the expression of key double-strand break repair effectors (16). Low RPS values indicate tumors with dysfunctional HR repair (16). *MYBL2* High tumors, regardless of tumor type, exhibited significantly elevated combined HRD scores and significantly decreased RPS scores, compared to *MYBL2* Low tumors (**Figure 2C**). Taken together, two orthogonal metrics indicate that *MYBL2* High tumors have inefficient HR repair.

Inefficient HR repair has been linked to inactivating mutations or deep deletions in BRCA genes (9). Given this, we profiled *MYBL2* High and Low tumors for somatic mutations or homozygous deletions in BRCA genes (**Figure 2C**). Surprisingly, mutations and deletions were rare in *MYBL2* High tumors. More importantly, these loss of function alterations were not significantly enriched when comparing *MYBL2* High and Low cohorts (**Figure 2C**). One exception to these findings was an enrichment for CHEK2 alterations in *MYBL2* High UCEC. Careful inspection of the data shown in Figure 2C reveals increased inactivating alterations in our LUAD and UCEC cohorts. This is likely because LUAD is linked to carcinogen exposure and several *MYBL2* High UCEC tumors carry POLE mutations, which impair polymerase proofreading. These results indicate that *MYBL2* High tumors fall into the clinically relevant category of tumors with genomic instability and inefficient HR repair despite carrying wildtype BRCA.

### Heterozygous loss of repair effectors underly defective DNA repair in *MYBL2* High tumors

Given the lack of BRCA gene inactivation, we characterized the DNA repair landscape in *MYBL2* High tumors in search of the genetic origin of genomic instability. To do this, we developed a weighted expression (WE) score to describe how expression of repair pathways is regulated in tumors (**Methods**). We applied this metric to all single-strand break repair pathways (SSBR), double-strand break repair pathways, and cell cycle checkpoint and lesion bypass mechanisms (**Figure 3A**). Analysis of WE pathway scores revealed a striking imbalance between expression patterns of different repair pathways. Key double-strand break repair pathways (HR, FA, MMEJ) and checkpoint signaling pathways were robustly upregulated across *MYBL2* High tumors while single-strand break pathways showed different degrees of downregulation (**Figure 3A**). Across tumor types, we found that translesion synthesis (TLS), nucleotide excision repair (NER), and non-homologous end-joining (NHEJ) pathways were consistently the most downregulated pathways. Correlation analysis demonstrated patterns observed in **Figure 3A** were strongly correlated across cancer types (**Figure 3B**). A notable tumor specific event was the strong downregulation of direct repair (DR) observed in *MYBL2* High IDH^MUT^ LGG.

**Figure 3:**
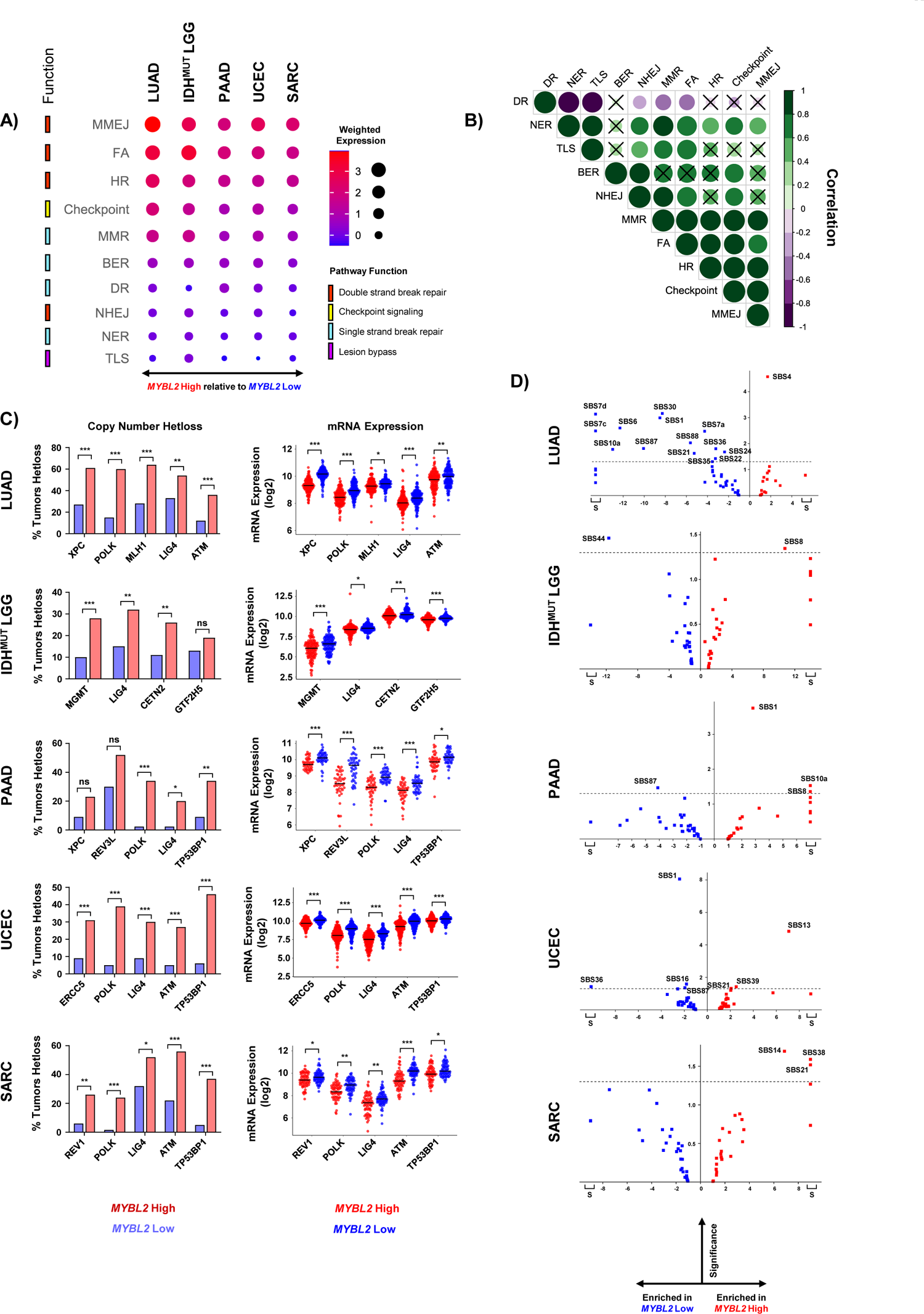
Heterozygous losses impacting key DNA repair effectors are enriched in *MYBL2* High tumors. **A)** Weighted expression scores reveal an imbalance in DNA repair pathway regulation. **B)** Observed differences in WE scores are highly correlated across different cancer types. Correlations with x marks indicate correlations that are not statistically significant (Pearson). **C)** Heterozygous losses in genes encoding key single-strand break repair, TLS, and NHEJ effectors are highly enriched in *MYBL2* High tumors. One-sided Fisher’s exact test, *p* < 0.05, *; *p* < 0.01, **; *p* < 0.001, ***. Heterozygous loss events are highly correlated with decreased expression of repair effectors. Benjamini-Hochberg corrected *q*. *q* < 0.05, *; *q* < 0.01, **, *q* < 0.001, ***. **D)** COSMIC v3.2 SBS analysis reveals heterozygous loss of repair effectors is associated with impaired pathway function. S: Signatures specifically observed only in *MYBL2* High or *MYBL2* Low tumors. Dotted line represents Student’s T-test p = 0.05.

We next asked if strongly downregulated pathway scores were predominantly driven by decreased expression of individual effector genes. Close inspection revealed that *MYBL2* High tumors exhibited strong downregulation of individual effectors (**Supplementary Table 3**). Using whole exome sequencing and copy number data, we profiled *MYBL2* High tumors for genetic alterations that could account for this specific downregulation. Like our BRCA gene analysis (**Figure 2C**), homozygous deletions and inactivating mutations were highly infrequent in *MYBL2* High tumors and could not explain the expression differences observed in **Figure 3A**. Additional analysis revealed that the driver of repair pathway dysregulation in *MYBL2* High tumors were specifically enriched heterozygous loss events impacting key repair effectors (**Figure 3C**). Importantly, these heterozygous loss events were highly correlated with decreased effector mRNA expression (**Figure 3C**). Looking across cancers, we found that heterozygous loss events in XPC (2/5 tumor types), POLK (4/5), LIG4 (5/5), ATM (3/5), and TP53BP1 (3/5) were common in *MYBL2* High tumors. Heterozygous loss of MGMT was specific to *MYBL2* High IDH^MUT^ LGG, fitting with previous reports of DR repair impairment being a tissue-specific driver of oncogenesis (17).

To assess the functional impact of these heterozygous loss events, we analyzed Catalog of Somatic Mutations in Cancer (COSMIC) v3.2 single-base substitution (SBS) signatures data (**Figure 3D**) (18). This analysis identified several SBS signatures that were enriched in *MYBL2* High tumors. For instance, SBS8 was over-represented in both *MYBL2* High IDH^MUT^ LGG and PAAD. SBS8 is characterized by increased C>A transversions and has been linked to deficient NER (18). This fits well given that *MYBL2* High IDH^MUT^ LGG have increased heterozygous losses in NER effectors CETN2 and GTF2H5 (**Figure 3C**). Also, *MYBL2* High PAAD have significantly increased heterozygous losses affecting both XPC and POLK. Here, XPC and POLK mediate the first (lesion recognition) and last (repair synthesis) steps of NER (19). SBS21 was over-represented in *MYBL2* High UCEC and SARC cohorts.

SBS21 is defined by increased T>C transversions and has been previously linked with NER defects (18). Previously, we identified that *MYBL2* High UCEC carried heterozygous losses in ERCC5 and POLK. Similarly, *MYBL2* High SARC also have significantly increased heterozygous losses in POLK. Lastly, signature SBS4 was over-represented in *MYBL2* High LUAD. SBS4 features increased C>A transversions and is the byproduct of tobacco-smoke induced lesions (20). Importantly, *MYBL2* High LUAD carried heterozygous losses in XPC and POLK which impair cellular ability to repair smoking induced lesions through NER (12). This analysis supports the notion that heterozygous losses of repair effectors functionally decrease pathway efficiency.

### Defective SSBR and TLS are linked to increased replication stress and distinct genomic footprints in *MYBL2* High tumors

SSBR and TLS pathways are essential for safe-guarding DNA replication. Various SSBR pathways are responsible for regulating the speed and accuracy of the replicative polymerases. Additionally, TLS represents an essential lesion bypass mechanism that helps alleviate replication fork stalling and collapse when the replicative machinery encounters DNA lesions (21). Genetic models of defective SSBR and TLS demonstrate significant genomic instability and elevated replication stress (22). Cells contain multiple pathways that sense and respond to replication dysregulation (23). Elevated expression of genes in these pathways are indicative of cells that experience significant replication stress (24). To investigate if *MYBL2* High tumors with impaired SSBR and TLS experience elevated replication stress, we developed a novel metric called the replication stress score (RS score) (**Methods**). This metric captures all major pathways involved in sensing replication stress, protecting and processing stalled replication forks, and the rescue of DNA replication (24). When comparing *MYBL2* High and Low cohorts, we found that *MYBL2* High tumors universally exhibited significantly elevated RS scores (**Figure 4A**). This suggests that *MYBL2* High tumors struggle with DNA replication, likely stemming from decreased SSBR and TLS capacity (**Figure 3**).

**Figure 4:**
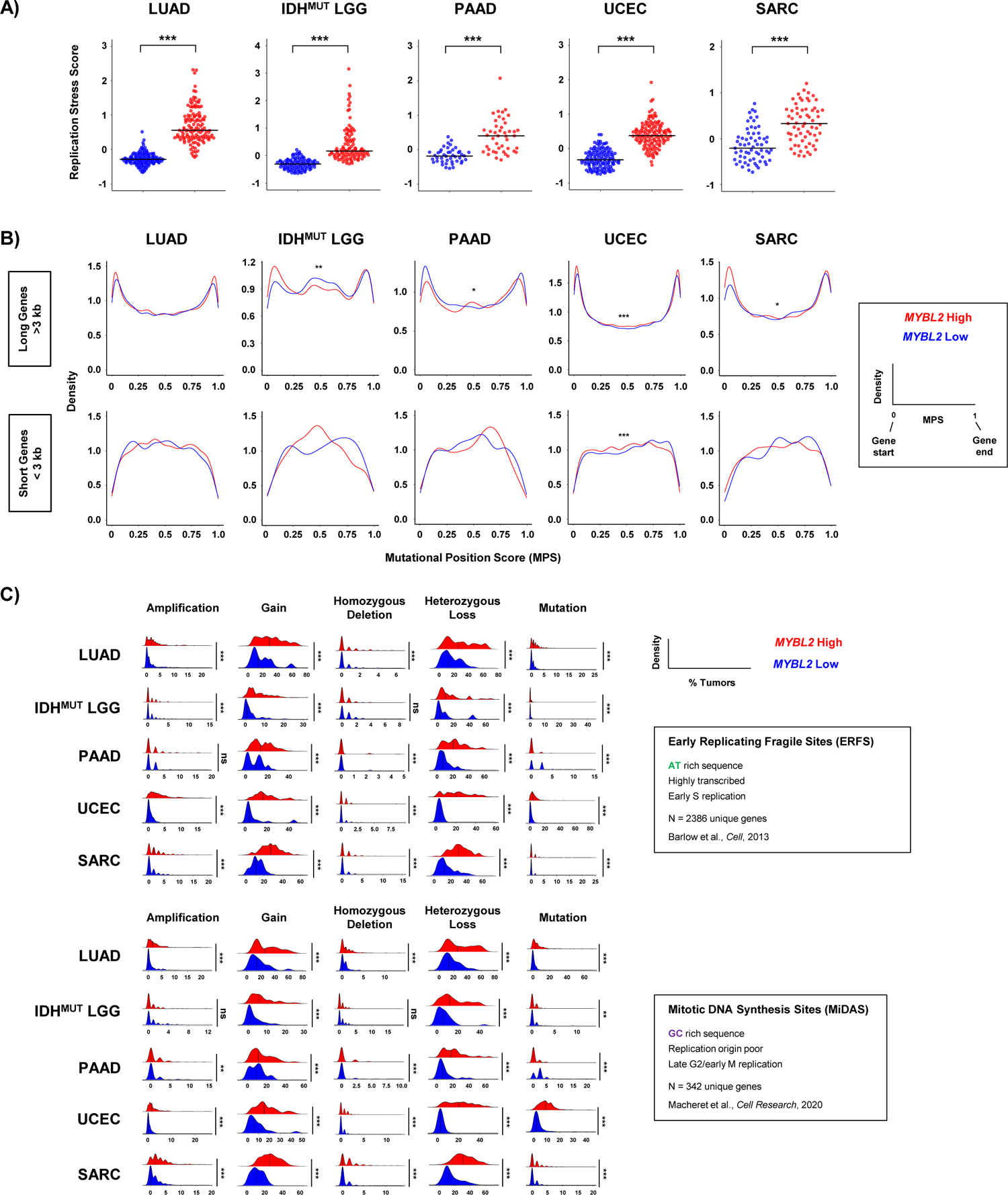
*MYBL2* High tumors exhibit markers of chronic replication stress. **A)** *MYBL2* High tumors universally demonstrate significantly elevated replication stress scores. Wilcoxon, *p* < 0.05, *; *p* < 0.01, **; *p* < 0.001, ***. **B)** *MYBL2* High tumors experience a shift in intragenic somatic mutation position, relative to *MYBL2* Low tumors. Kolmogorov-Smirnov test, *p* < 0.05, *; *p* < 0.01, **; *p* < 0.001, ***. **C)** *MYBL2* High tumors acquire significantly greater numbers of alterations at replication stress sensitive genomic sites. Wilcoxon, *p* < 0.05, *; *p* < 0.01, **; *p* < 0.001, ***.

Based on findings in Figure 4A, we next asked if *MYBL2* High tumors accumulate somatic mutations at different locations and frequencies across intragenic regions. To test this hypothesis, we developed a metric called the mutational position score (MPS) (**Methods**). This metric allows us to directly compare the spatial location of somatic mutations in individual genes across all tumors. Here, MPS values closer to 0 correspond to mutations near to the gene start, while values near 1 correspond to mutations close to the gene end. MPS values near 0.5 represent intragenic mutations accumulating in the middle of the gene body. When comparing MPS density traces, we found that *MYBL2* High tumors experience significant shifts in intragenic mutation location frequency (**Figure 4B**). For this analysis, we subdivided all genes based on gene length into long genes (> 3000 bp) and short genes (<3000 bp). For long genes, *MYBL2* High tumors tended to acquire more mutations near gene starts and gene ends, likely stemming from transcription-replication conflicts. *MYBL2* High PAAD tumors, however, interestingly showed increased accumulation of mutations near the middle of long genes. Analysis of short genes showed even more pronounced changes, where *MYBL2* High tumors showed increased mutation in the body of short genes (**Figure 4B**). The lack of significance for these patterns likely stems from fewer mutations in short genes, compared to long genes. Collectively, this shift in mutational position is consistent with increased replication stress and impaired SSBR and TLS pathways seen across *MYBL2* High tumors.

It has long been understood that thousands of genomic loci are sensitive to replication stress (25). Recent studies have subdivided these loci into two categories, early replicating fragile sites (ERFS) and mitotic DNA synthesis sites (MiDAS) (26–27). Genes encoded at these sites are sensitive to replication stress due to their local DNA sequence, replication timing, and location in the genome. ERFS genes have been shown to be highly AT rich, highly transcribed, and replicated in early S phase (26). As a result, the replicative polymerase frequently slips or encounters an RNA-polymerase, causing stalling or DNA breaks. These events have been shown to cause early replicating sites to be gained or amplified at increased rates. MiDAS genes, on the other hand, contain highly GC rich sequences, are replicated in late G2/M, and are located in replication origin poor regions (27). These circumstances make MiDAS genes difficult to replicate and cells frequently commit to mitosis prior to completing replication at these sites. Late replicating genomic regions have been associated with increased deletions as cells use various methods to complete replication (28). Given this, we hypothesized that *MYBL2* High tumors acquire greater numbers of genomic alterations at replication stress sensitive (RSS) sites. Using copy number and WES data, we profiled *MYBL2* High and Low tumors for amplifications, gains, homozygous deletions, heterozygous losses, and mutations impacting ERFS and MiDAS sites (**Methods**). Across both ERFS and MiDAS loci, we found that *MYBL2* High tumors accumulate significantly greater numbers of genetic alterations (**Figure 4C**). Strikingly, we found that the number of gene-level gains and heterozygous losses dwarfed that observed for amplifications, homozygous deletions, or mutations. Additionally, copy number trends associated with replication timing did not correlate with our findings (28); *MYBL2* High tumors acquired similar numbers of gains and heterozygous losses across both ERFS and MiDAS loci, with a trend toward more heterozygous losses (**Figure 4C**). This data is consistent with previous findings where elevated MMEJ activity is coincident with increased loss of heterozygosity events (29). Across all tumor types, we observed strong right-handed tailing indicating that many genes are impacted by gains or heterozygous losses in greater than 30-40% of *MYBL2* High tumors (**Figure 4C**). Taken together, these data suggest that repeated gene-level gains and heterozygous losses at RSS genomic sites originate from increased replication stress.

### Recurrent copy number alterations at RSS sites rewire transcriptional programs and impact hallmark of cancer master regulators

After noticing that large numbers of genes were recurrently altered in *MYBL2* High tumors, we examined the function of genes encoded at RSS genomic sites (**Methods**). Biological process analysis revealed that genes encoded at RSS sites fit into thirteen functional categories (**Figure 5A**). Importantly, we found that conserved copy number changes significantly impacted gene expression (**Figure 5A-B**). Across cancers, we found that *MYBL2* High tumors frequently gained copies of genes controlling DNA replication and repair (cluster 4). Similarly, we found recurrent heterozygous losses impacting multiple genes controlling cell death and survival (cluster 7). While some events were confined to individual tumor types, there was striking conservation of both the number and identity of genes altered across functional clusters in *MYBL2* High tumors (**Figure 5A, Supplementary Figures 1-5**). Further analysis revealed that copy number alteration and subsequent transcriptional regulation impacted master effectors responsible for regulating several hallmarks of cancer. Specifically, we observed repeated heterozygous loss and transcriptional downregulation of *TMEM173/STING1* (evading immune surveillance), *DAPK2* (evading cell death), *POLK* (DNA damage), *JAK2* (evading immune surveillance), *NF1* (growth factor signaling), *PDCD4* (protein translation), and *MGMT* (DNA damage) (**Figure 5B**). Recurrent copy number gains and transcriptional upregulation was observed for *BCL2L1* (evading cell death), *LIN9* (transcription), *ZEB1* (cell movement), MYC (transcription), *TK1* (limitless replicative potential), *LIN37* (transcription), and *ERBB2/HER2* (growth factor signaling) (**Figure 5B**). These results indicate that increased replication stress, stemming from heterozygous repair effector loss, promotes dysregulation of key master regulators which are encoded at RSS sites (**Figure 5C**).

**Figure 5:**
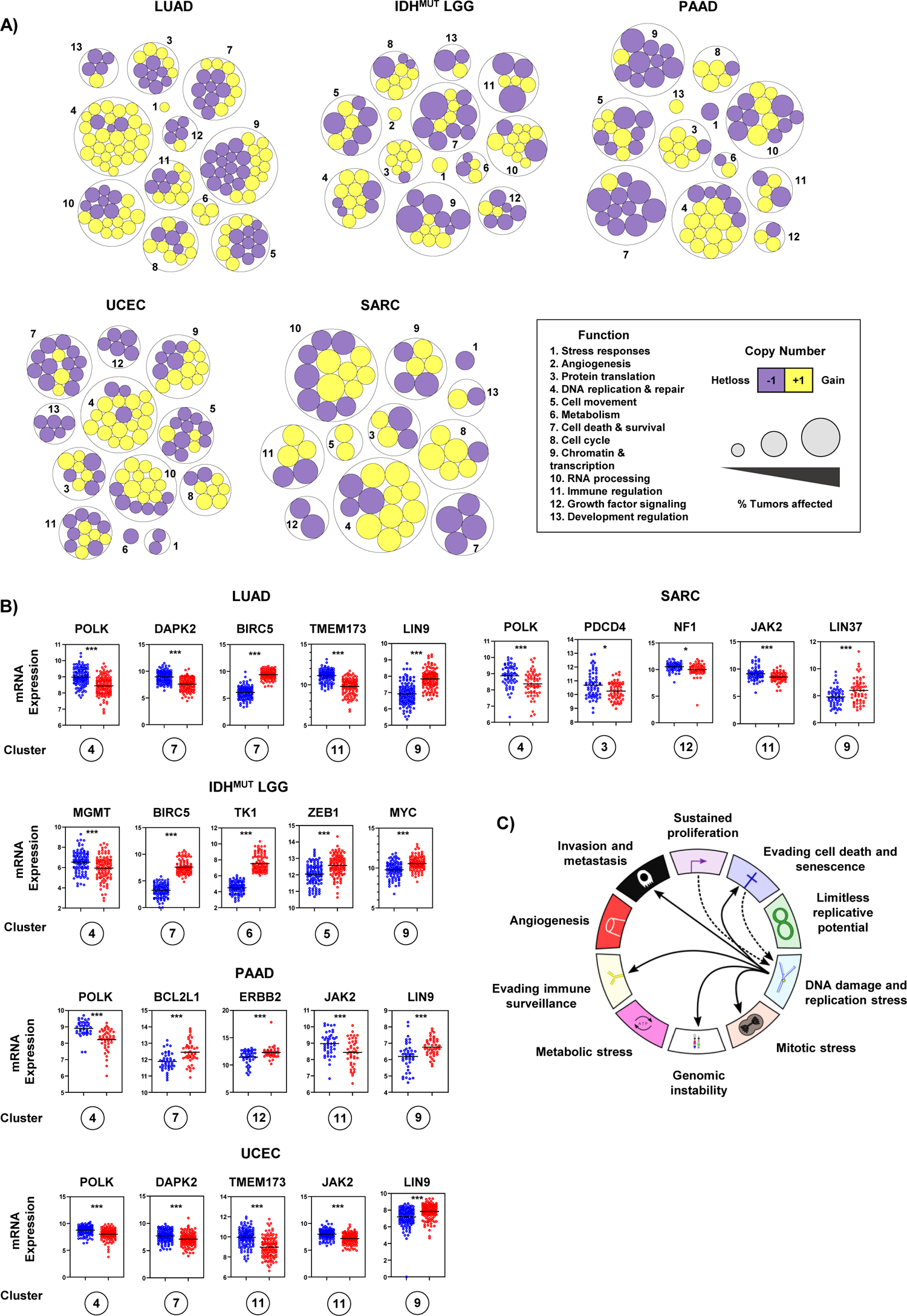
Recurrent copy number alterations at replication stress sensitive sites rewire transcriptional programs and dysregulate master effectors controlling several hallmarks of cancer. **A)** *MYBL2* High tumors acquire copy number alterations in essential enzymes encoded at replication stress sensitive sites. **B)** Enriched copy number alterations observed in *MYBL2* High tumors rewire transcriptional programs and dysregulate master effectors controlling several hallmarks of cancer. Statistical significance is mapped according to Benjamini-Hochberg corrected *q* values. *q* < 0.05, *; *q* < 0.01, **; *q* < 0.001, ***. Circled cluster numbers map to those displayed in **A)**. **C)** Replication stress dysregulates master effectors controlling several hallmarks of cancer.

### *MYBL2* High tumors exhibit increased neoantigen loads and immunosuppressive microenvironments

Given an increased dysregulation of key effectors controlling immune regulation, we sought to characterize the immune microenvironment associated with *MYBL2* High tumors. As expected, we found that *MYBL2* High tumors have significantly greater neoantigen loads compared to *MYBL2* Low (**Figure 6A**) (30). Next, we used ConsensusTME and TIDE algorithms to generate infiltration estimates for immune and stromal cell subtypes (31–32). Interestingly, we found that *MYBL2* High tumors across tumor types lacked statistically significant differences in CD8+ T-cell infiltration (**Figure 6B**). However, *MYBL2* High tumors universally were associated with elevated infiltration of myeloid-derived suppressor cell (MDSC) populations (**Figure 6B**). Across tumor types, we also found that *MYBL2* High tumors were associated with greater Exclusion scores and decreased Dysfunction scores (**Figure 6B**). One exception to this trend was *MYBL2* High PAAD, where both measures were trending but not statistically significant, likely due to smaller patient cohort sizes. Lastly, analysis of tumor hypoxia scores revealed *MYBL2* High tumors are significantly hypoxic (**Figure 6C, Supplementary Figure 11**) (13). Increased hypoxia scores fit well with increased MDSC infiltration estimates and significantly decreased infiltration of endothelial cells across *MYBL2* High tumors (**Supplementary Figures 6-10**). Hypoxia scores for TCGA SARC tumor samples were not available and are not included in the analysis in **Figure 6C**. All together, these data indicate that despite harboring increased neoantigen loads, *MYBL2* High tumors exhibit uniquely dysregulated, immunosuppressive microenvironments.

**Figure 6:**
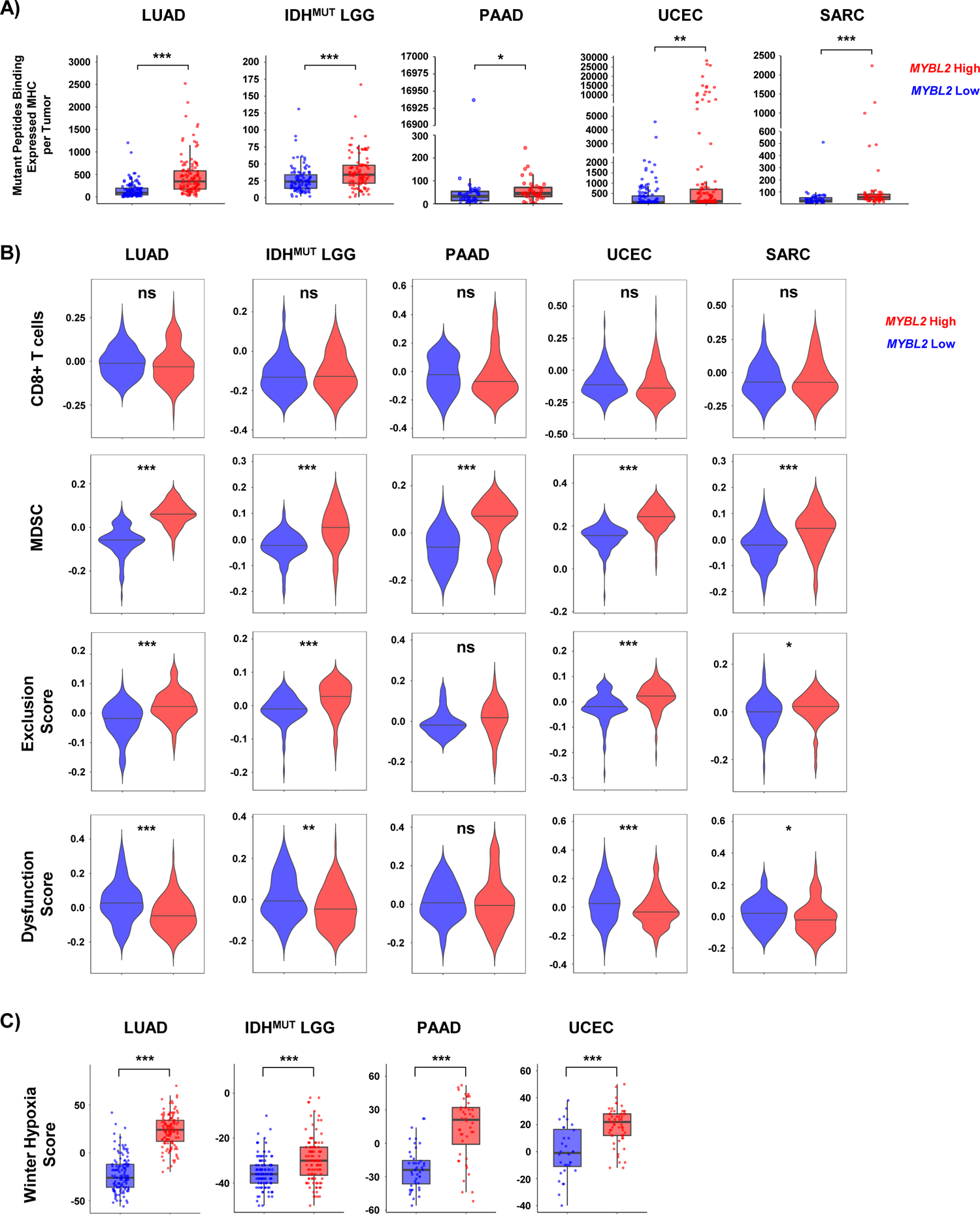
*MYBL2* High tumors exhibit uniquely dysregulated tumor microenvironments. **A)** *MYBL2* High tumors contain significantly greater numbers of mutant peptides that bind to patient-matched, expressed, pMHC complexes. **B)** Immune infiltration estimation algorithms indicate that *MYBL2* High tumors are significantly more immunosuppressive. **C)** *MYBL2* High tumors are highly hypoxic. **A), B), C)** Wilcoxon, *p* < 0.05, *; *p* < 0.01, **; *p* < 0.001, ***.

### Elevated *MYBL2* expression identifies patients at increased risk for therapy failure and distant metastases

Next, we sought to investigate the association of this *MYBL2* High phenotype with therapy response. To test if elevated *MYBL2* expression identified patients with poor responses to therapy, we analyzed 25 tumor types provided by the Oncology Research Information Exchange Network (ORIEN). Kaplan-Meier analysis demonstrated that patients with *MYBL2* High tumors had significantly poorer overall survival outcomes when treated with chemotherapeutics or irradiation across LUAD, IDH^MUT^ LGG, invasive ductal breast cancer (ID-BRE), and late-relapse multiple myeloma (LRMM) cohorts (**Figure 7A**). These results fit well with our TCGA analyses where we linked elevated *MYBL2* expression with poor outcomes in treatment naïve LUAD and IDH^MUT^ LGG (**Figure 1**). For ID-BRE, elevated *MYBL2* expression was not prognostic in our TCGA analysis, despite showing a similar biology to that of other *MYBL2* High cohorts described throughout this study (**Supplementary Table 1**). However, elevated *MYBL2* expression was highly predictive when patients were treated with chemotherapeutic or irradiation regimens. This analysis also extended our results into liquid tumors with *MYBL2* expression being robustly prognostic in the most recalcitrant form of multiple myeloma, LRMM (>4 lines of prior therapy). Importantly, analysis of COSMIC SBS v3.2 signatures confirmed resistant *MYBL2* High tumors demonstrate footprints of defective SSBR and TLS effector function (**Supplementary Figure 12**). We also developed FUSED to nominate error-prone repair pathways responsible for generating genomic fusions detected by RNA-seq (**Methods**). In *MYBL2* High samples that responded poorly to therapy, we found evidence of elevated MMEJ activity (**Supplementary Figures 13-16**). Collectively, these results demonstrate that DNA repair defects and increased error-prone repair potentiating *MYBL2* High disease is linked to poor responses to chemotherapy and irradiation across tumor types.

**Figure 7:**
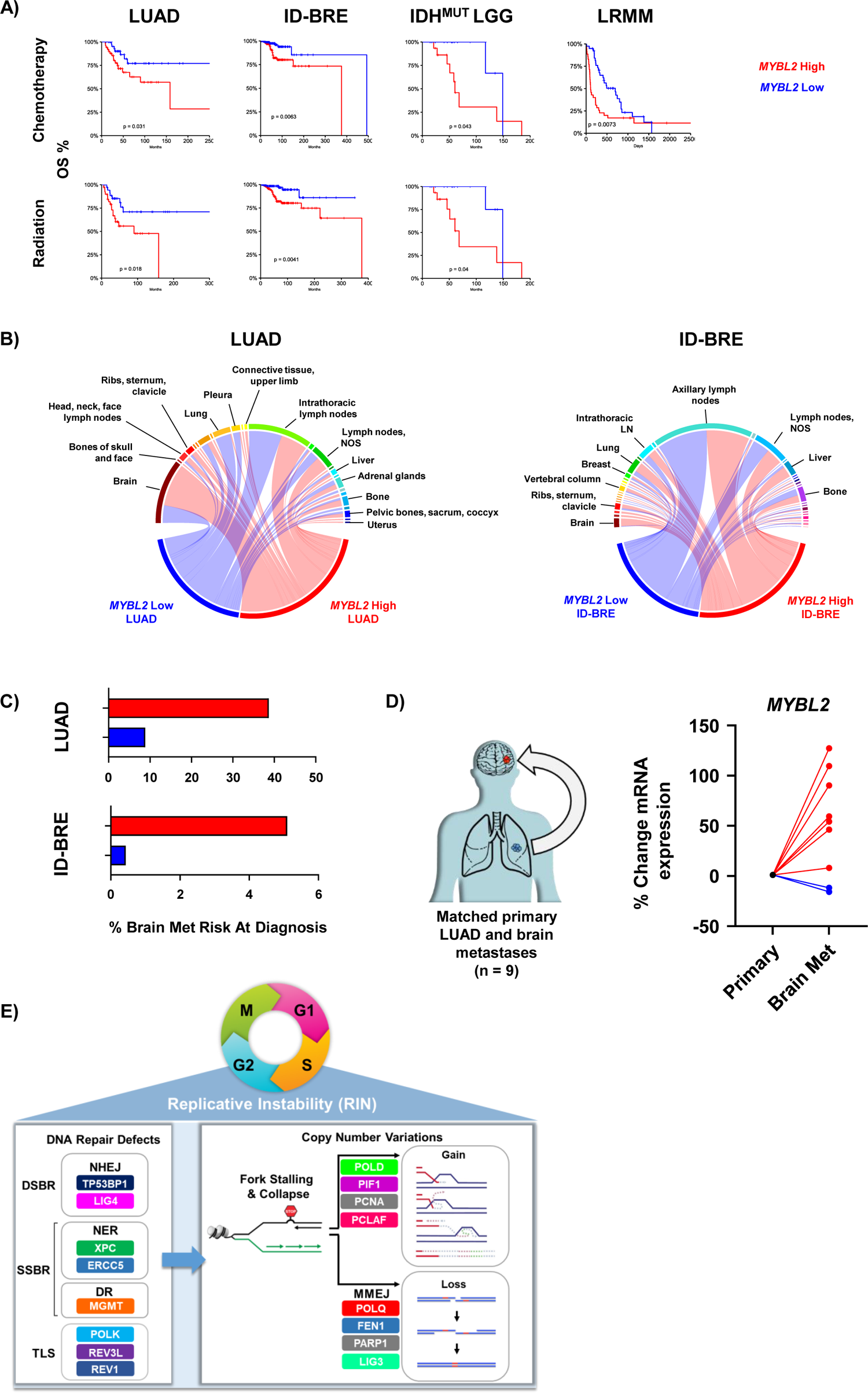
Elevated *MYBL2* expression identifies patients at risk for poor responses to therapy and distant metastases across tumor types. A) *MYBL2* High patients have significantly poorer outcomes when treated with chemotherapy and irradiation regimens. Log-rank test *p*-values are displayed. B) *MYBL2* High tumors metastasize to distant sites at a higher frequency, including to the brain. C) *MYBL2* expression stratifies patient risk at diagnosis for brain metastasis development. LUAD: lung adenocarcinoma. ID-BRE: Invasive ductal breast cancer. IDH^MUT^ LGG: IDH-mutant lower grade glioma. LRMM: Late relapse multiple myeloma. D) *MYBL2* expression is increased in brain metastases compared to patient matched primary lung adenocarcinoma tumors. E) Replicative instability (RIN) accelerates genome evolution, driving cancer progression.

Lastly, we analyzed patient records to assess for potential differences in metastatic dissemination (**Methods**). When comparing *MYBL2* High and Low cohorts, we found no difference in dissemination to sentinel lymph nodes in both LUAD (intra-thoracic lymph nodes) and ID-BRE (axillary lymph nodes) cohorts (**Figure 7B**). However, we found that *MYBL2* High tumors demonstrated increased dissemination to distant metastatic sites in both LUAD and ID-BRE, especially to the brain. Interestingly, we found no difference in the median time to metastasis between *MYBL2* High and Low cohorts, suggesting that observed patterns reflect tissue-specific tropisms (**Supplementary Figure 17**). Using combined probability, we found that *MYBL2* expression dramatically stratifies patient risk at diagnosis for developing brain metastases during their disease course (**Methods, Figure 7C**). Importantly, these values match or exceed current genomic markers for brain metastasis risk for both LUAD and ID-BRE (33). Strikingly, analysis of primary lung adenocarcinoma and paired brain metastasis samples revealed that *MYBL2* expression significantly increased in 7 of 9 samples (**Figure 7D**). These data suggest that *MYBL2* may be a putative driver of primary carcinoma to brain metastatic dissemination.

## Discussion

Across multiple tumor types, elevated *MYBL2* expression identified tumors with genomic instability, inefficient homologous recombination, and wildtype BRCA (**Figures 1-2**). Analysis of the DNA repair landscape revealed that the genetic basis of *MYBL2* High disease are heterozygous losses of SSBR, TLS, or NHEJ effectors (**Figure 3**). We found that these heterozygous losses were linked to elevated replication stress, a shift in intragenic mutation position, and increased copy number alterations in genes encoded at RSS genomic sites (**Figure 4**). Functional clustering approaches allowed us to discover that replication stress promotes copy number alterations that rewire transcriptional programs regulating hallmarks of cancer master effectors (**Figure 5**). Clinically, this phenotype identifies patients at risk for poor responses to chemotherapy and irradiation (**Figure 7**). Additionally, our results demonstrate that patients with *MYBL2* High disease are at increased risk for distant metastases, especially to the brain (**Figure 7**).

In this study, we have identified a new cohort of tumors characterized by replicative instability (RIN) (**Figure 7E**). Across multiple tumor types, we find that RIN tumors exhibit significant FGA, increased MSI, and elevated somatic mutations (**Figure 2**). At the chromosomal level, RIN tumors demonstrate significantly greater levels of intrachromosomal alterations, such as gene-level gains and heterozygous losses, likely caused by stalled or collapsed DNA replication intermediates (**Figure 2, Figure 4**). Importantly, we found that RIN is coincident with heterozygous losses of key SSBR, TLS, and NHEJ repair effectors (**Figure 3**). As a consequence, RIN tumors upregulate genes controlling the replication stress response, MMEJ, FA, and checkpoint machinery (**Figure 3**). Unlike chromosomal instability (CIN), our work supports a model in which RIN accelerates genomic evolution during replication, as opposed to missegregation during mitosis. It is important to note that RIN develops across cancer genotypes and tissue types. Analysis of tissue-specific driver events revealed that RIN was not consistently linked to specific driver alterations (**Figure 1**).

Additionally, we find that RIN develops across cancers in the lung, brain, pancreas, uterus, connective tissue, breast, and hematopoietic compartment (**Figure 1, Figure 7**). This phenotype likely extends to other tumor types besides those described here (**Supplementary Table 1**). In this manuscript, we show that elevated *MYBL2* expression and RIN are intimately linked. As described below, the association of MYBL2 with RIN is both direct and indirect. In normal cells, MYBL2 is transcriptionally and post-translationally regulated by the cell cycle (34). Specifically, MYBL2 is transcriptionally upregulated as cells enter S-phase. During S-phase, MYBL2 is phosphorylated by CCNA:CDK2 and actively regulates transcription. As cells progress through G2, MYBL2 upregulates the expression of FOXM1 and other effectors that promote G2/M progression. MYBL2 is then hyper-phosphorylated by CCNA:CDK2 and targeted for degradation to allow cell division. Given this, elevated expression of *MYBL2* mRNA is a robust marker of cells that are arrested prior to mitosis. In this study and our previous work, we have demonstrated that *MYBL2* expression is tightly associated with the transcriptional upregulation of DNA repair genes that sense replication stress (12). This fits well when considering the mechanisms through which these signaling pathways coordinate cell cycle arrest following replication stress. Upon replication stress, ATR activates its effector kinase, CHK1 (23). CHK1 then phosphorylates CDC25 family members and inhibits their phosphatase activity, halting cell cycle progression. By doing so, CHK1 prevents CCNB1:CDK1 activity that prevents cells from progressing to mitosis. Importantly, increased MYBL2 expression and transcriptional activity are indirect effects of CHK1 mediated cell cycle arrest. This indicates that increased MYBL2 expression and activity promotes genomic evolution during replication, driving RIN. Taken together, increased *MYBL2* expression and transcriptional activity are robust markers of RIN.

Therapy resistance and metastasis are key cancer progression events that directly impact survival outcomes. Our results indicate that RIN tumors respond poorly to chemotherapy and irradiation. These findings fit well when considering the genetic background of these tumors. Chemotherapy and irradiation regimens are designed to overwhelm the replicative machinery, causing cell death. Several studies have demonstrated that upregulation of inter-strand crosslink repair (FA), cell cycle checkpoint signaling, and error-prone repair (MMEJ) pathways confer resistance to these therapies (35–36). Because RIN tumors carry heterozygous losses in key SSBR, TLS, and NHEJ effectors, they experience chronic replication stress. To cope with this stress, therapy naïve tumors upregulate FA, cell cycle checkpoint, and MMEJ pathways. In doing so, these tumors become primed for resistance to DNA damaging therapies. In addition to therapy resistance, heightened replication stress and elevated error-prone repair pathway activity promote copy number alterations in key regulators of hallmarks of cancer processes. For instance, we find that this mechanism underlies dysregulation of TMEM173, JAK2, DAPK2, BIRC5, LIN9, LIN37, ERBB2, and NF1, among others (**Figure 5**). Dysregulation of these and other crucial effectors allow cancers to evade the immune system, resist anoikis driven apoptosis, achieve growth-factor independent signaling, and move. Additionally, gains in LIN9 and LIN37 further potentiate this phenotype by increasing MYBL2 expression and transcriptional activity. These alterations dramatically shorten the molecular time required for developing an aggressive cancer capable of distant metastases. Consistent with this, we find that *MYBL2* High LUAD and ID-BRE tumors are more likely to metastasize to distant sites, especially to the brain. Collectively, our results indicate that RIN is a pan-cancer driver of progressive disease.

Our results have important implications for treatment plans and clinical trial design. As RIN tumors respond poorly to chemotherapy and irradiation, clinical trials should explore targeted therapy combinations in the therapy refractory setting. Given that RIN tumors display large quantities of neoantigens, the question of immunotherapy response is highly relevant. Because therapy naïve RIN tumors exhibit highly hypoxic, MDSC-rich microenvironments, it may be unlikely that these tumors achieve durable responses to anti-PD1/PDL1 inhibitors. However, in our ORIEN cohorts, we find that RIN tumors are associated with increased LAG3 and TIGIT expression, despite showing no difference in PDL1 (data not shown). This raises the possibility that new anti-LAG3 and anti-TIGIT immune checkpoint inhibitors may be better suited for treating RIN tumors. One of our most important discoveries is that increased *MYBL2* expression, and thus RIN, dramatically stratifies patient risk for brain metastases in LUAD and ID-BRE. While the average risk for brain metastases for all lung cancers is reported to be 15%, we find that *MYBL2* High patients have a risk of ∼40% while *MYBL2* Low have a risk of less than 10% (**Figure 7**) (50). A similar dichotomy is observed in ID-BRE, where the reported risk for brain metastases for breast cancer patients is ∼5%. Here, we find that *MYBL2* High ID-BRE risk is 5% while *MYBL2* Low ID-BRE is <1%. These results strongly argue for increased screening for brain metastases in patients with *MYBL2* High disease.

Moving forward, further study of RIN is urgently needed. Given the aggressive nature of RIN tumors, immunohistochemistry markers need to be identified and validated. New mouse models and cell line systems are required in order identify potential therapeutic vulnerabilities that can be explored in clinical trials. Any advances in identifying and targeting RIN have the potential to drastically improve patient outcomes across multiple tumor types.

## Methods

### TCGA pan-cancer analysis

Thirty-two tumor types curated by the TCGA and other groups were analyzed in this study (13). Where multiple TCGA studies were available, we focused our analyses on PanCancer studies. Samples with RNA-sequencing data were stratified into *MYBL2* High and *MYBL2* Low cohorts using normalized mRNA expression values and a quartile method; the top 25% of samples expressing *MYBL2* mRNA were called *MYBL2* High and the bottom 25% of samples *MYBL2* Low.

### Survival Analyses

The Kaplan-Meier estimator was used to estimate time-to-event distributions for OS, DSS, and PFS outcomes. The log-rank test was used to test for significant differences between distributions using a two-sided test. OS denotes the time from initial diagnosis until death. DSS is defined as the time from cancer diagnosis until the time of death; patients who died from other causes were not included. PFS reflects the time from initial diagnosis until progression or death. For all three survival analyses, patients who did not experience an event or who were lost to follow-up were censored at the time of last contact. Kaplan-Meier survival analyses were conducted using survival and survminer R packages (37).

### DNA repair pathway WE score

A WE score was developed to describe how DNA repair pathways are regulated in tumors. For each repair pathway, we identified comprehensive lists of pathway effectors through extensive literature review (19, 21, 23, 38–44). Effectors were scored based on essentiality to pathway function (Essentiality Scaling Factor: 3 = essential effector, 2 = important effector or potentially compensable, 1 = accessory effector). The final WE formula for each pathway is a scaled average where gene mRNA Log2FC values are multiplied by an essentiality scaling factor (ESF), summed, and divided by the number of pathway genes. Correlations between WE values were calculated and visualized using stats and corrplot R packages (45).

### COSMIC v3.2 SBS analysis

COSMIC SBS v3.2 signatures were generated using the deConstructSigs R package (46). For TCGA cohorts, the TCGA public MAF file (mc3.v0.2.8.PUBLIC.maf.gz) was used to generate trinucleotide mutation context matrices. For ORIEN cohorts, individual sample vcf files were used to calculate trinucleotide mutation contexts. The final deConstructSigs output was computed using the trinucleotide context matrix and the COSMIC v3.2 SBS mutational signature matrix downloaded from (https://cancer.sanger.ac.uk/signatures/downloads/). Eighteen of the 78 SBS mutational signatures likely capturing sequence artifacts were excluded. Statistical significance between average signature weights across samples was assessed using two-sided Student’s T-tests.

### Replication stress score

To analyze differences in replication stress, we developed the replication stress (RS) score. Eight gene ontology (GO) terms were identified that capture key cellular processes involved in replication stress responses (GO:0031570, GO: 0000076, GO: 006260, GO: 0031261, GO: 004311, GO: 0031297, GO: 0031298, GO: 0071932). Genes were pooled and redundant entries removed to generate a final gene list (n = 205). The RS score is the sum of gene log2 mRNA expression values, divided by the total number of genes in the RS response gene list. Differences in medians were assessed for statistical significance using Wilcoxon signed rank tests (47).

### Mutational position score

The mutational position score (MPS) was developed to assess differences in intragenic mutation frequency. Here, the MPS score is the difference between somatic mutation location and the gene start, divided by gene length. Mutation locations were obtained from the TCGA public MAF file. Gene start and end positions were obtained from Ensembl. Differences in mutational position densities were assessed for statistical significance using Kolmogorov-Smirnov tests.

### RSS genomic site alteration analysis

RSS genomic sites were identified by Barlow et al. (26) and Macheret et al. (27). ERFS were obtained from Table S1 “Ordered_List_of_ERFS_Hot_Spots” (26). These genes were mapped to human gene identifiers using the nichenetr R package (48). A list of MiDAS sites was obtained from Supplementary Table S1 (27). Sites were filtered to include MiDAS sites attributable to one or two genes (removes unmappable intergenic sites). Final ERFS and MiDAS sites were merged to identify any overlapping genes. This merge identified 20 genes identified as ERFS but recently defined as MiDAS sites. These genes were subsequently removed from the ERFS list and only analyzed in the MiDAS list. Copy number alteration and somatic mutation frequencies were plotted using ggplot2 and ggridges R packages. Differences in medians were assessed for statistical significance using Wilcoxon signed rank tests (47).

### RSS site functional analysis

ERFS and MiDAS genes were combined into a single list and analyzed for broad biologic processes using WebGestalt’s over-representation analysis feature. From this analysis, thirteen functional clusters were defined and genes were binned into clusters following literature review (**Supplementary Table 5**). Single-cell RNA expression data from the Human Protein Atlas was used to ensure genes were expressed in tissues relevant to our tumor cohorts. This final gene list with functional cluster annotation was then merged with differential expression RNA-seq tables. Combined copy number and RNA-seq expression files were analyzed and genes with significant copy number and transcriptional differences were identified (**Supplementary Table 6**). Circular packing diagrams were drawn using ggraph and igraph R packages (49).

### Tumor microenvironment analysis

Immune cell infiltration estimates were generated using RSEM gene normalized values and the ConsensusTME R package (31). Individual tumor type infiltration estimates were calculated separately using tumor specific gene sets and a ssgsea method. Myeloid derived suppressor cell (MDSC) infiltration estimates, Dysfunction, and Exclusion Scores, were downloaded from the TIDE database (32).

### ORIEN therapy response analysis

Data from 25 tumor types provided by ORIEN in the May 2021 private cBioPortal instance were analyzed. For samples with RNA-seq data, we manually reviewed treatment records to identify patients treated with chemotherapeutics and/or irradiation. For treatment specific cohorts, we used normalized RNA expression values to stratify patients into *MYBL2* High and *MYBL2* Low cohorts using a quartile method. Kaplan-Meier analyses were performed as described above.

#### FUSED

<u>FUS</u>ion <u>E</u>rror-prone repair <u>D</u>etection (FUSED) was developed to map the origin of RNA-seq detected fusions. FUSED identifies fusions with closest similarity to NHEJ, single strand annealing (SSA), MMEJ, break induced replication (BIR), or microhomology mediated break induced replication (MMBIR). Tool rules were determined through literature review (51). FUSED is publicly available, https://github.com/databio/FUSED.

### ORIEN metastatic dissemination analysis

ORIEN medical records were manually reviewed to identify sites of metastatic disease. Metastatic dissemination routes were plotted using the circlize R package (50). Medical records were used to calculate the time from diagnosis to metastatic disease development. Time to metastatic disease distributions were plotted using the swimplot R package. Differences in time to metastasis data were assessed using Wilcoxon signed rank tests (47). Brain metastasis risk was calculated by multiplying the number of patients that develop metastatic disease by the number of patients with brain metastases. This fraction was multiplied by 100% to generate the final risk percentage.

### Statistical analyses

Statistical tests for all analyses are indicated in accompanying figure legends. For all boxplots, data are graphed as minimum, 1^st^ quartile, median, 3^rd^, quartile, and maximum. *p* and *q* values < 0.05 were considered statistically significant.

## Acknowledgements

The authors would like to acknowledge the following ORIEN member institutions for their commitment to data sharing and for contributing samples to this study: the University of Virginia Comprehensive Cancer Center, Moffitt Cancer Center, The Ohio State University Comprehensive Cancer Center, Markey Cancer Center, Huntsman Cancer Institute, University of Southern California Norris Comprehensive Cancer Center, Indiana University Melvin and Bren Simon Comprehensive Cancer Center, University of Iowa Holden Comprehensive Cancer Center, Roswell Park Comprehensive Cancer Center, University of Colorado Comprehensive Cancer Center, Emory Winship Cancer Institute, Rutgers Cancer Institute of New Jersey, and Stephenson Cancer Center. ORIEN molecular data analyzed in this study were managed by M2Gen under the Total Cancer Care (TCC) protocol at ORIEN member institutions. The authors also acknowledge the contributions of each institution’s ORIEN Team and biorepository and tissue research facility staff in the consent of patients, specimen procurement, specimen processing, data abstraction, and providing access to molecular and clinical data (UVA IRB HSR 18445, HCI IRB #89989, Moffitt Advarra IRB Pro00014441, Markey IRB #44224, Emory Winship IRB # 00095411, USC Norris HS-16-00050, Stephenson Cancer Center IRB #8323, University of Iowa Holden IRBs #201708847 and 201003791, University of Colorado Comprehensive Cancer Center IRB # 15-1110, Indiana University Melvin and Bren Simon Comprehensive Cancer Center IRB # 1807389306, and Roswell Park Comprehensive Cancer Center IRB # I 03103.). We are also grateful for the expert assistance and service of the PKPD Core at Moffitt Cancer Center as well as the members of the Pentecost Family Myeloma Research Center (PRMC) at Moffitt. The authors thank Gabriel De Avilla and Raghu Reddy Alugubelli for their help in multiple myeloma sample processing. The authors sincerely thank our patients and their families for donating their samples for research purposes. The authors would like to thank Dr. Yuh-Hwa Wang and Heather Raimer-Young for their insightful discussions and comments.

## Author Contributions

BBM conceptualized the study, conducted formal analysis, visualized data, developed software, wrote the original draft, reviewed, and edited the manuscript, and acquired funding to support this study. JPS conducted formal analysis, developed software, and reviewed and edited the final submission. QZ, ZJ, OAH, and MLC performed formal analysis, curated data, conducted formal analysis, data visualization, and helped review and edit the final manuscript. SMA, DHO, JEG, PMD, HHS, DGS, HC, AC, MV, KHS, AS, JLV, VFB, WLA, RDG, RDH, CBM, CMU, ARP, DAN, EAS and JML provided resources and wrote, edited, and reviewed the manuscript. DRJ and PTS wrote, reviewed, and edited the manuscript. MWM conceptualized the study, acquired funding, supervised the study, and wrote, reviewed, and edited the manuscript. All authors contributed to the article and approved the submitted version.

## Data availability statement

Some data analyzed in this study are subject to the following licenses/restrictions: Access to ORIEN data is controlled by M2Gen and the ORIEN consortium. Requests to access these datasets should be directed to https://www.oriencancer.org/request-an-account. Publicly available data sets were analyzed in this study. Tumor type specific survival, clinical, and genomic data can be found in cBioPortal (https://www.cbioportal.org/) under the following studies: Lung Adenocarcinoma (TCGA, PanCancer Atlas), Brain Lower Grade Glioma (TCGA, PanCancer Atlas), Pancreatic Adenocarcinoma (TCGA, PanCancer Atlas), Uterine Corpus Endometrial Carcinoma (TCGA, PanCancer Atlas), and Sarcoma (TCGA, PanCancer Atlas). Mutation Position Scores were generated using TCGA MAF file mc3.v0.2.8.PUBLIC.maf.gz (gdc.cancer.gov/about-data/publications/pancanatlas). Copy number analyses for MiDAS and ERFS genes were conducted using SCNV gene level, GITSTIC 2 thresholded files for LUAD, LGG, PAAD, UCEC, and SARC studies (linkedomics.org). ConsensusTME scores were generated using RSEM gene normalized RNA-seq files downloaded from GDAC FireBrowse (firebrowse.org) for each tumor study. Neoantigen and pMHC data are available from Thorsson et al. (Supplementary files: TCGA_PCA.mc3.v0.2.8.CONTROLLED.filtered.sample_neoantigens.10062017.tsv, TCGA_pMHC_SNV_sampleSummary_MC3_v0.2.8.CONTROLLED.170404.tsv, gdc.cancer.gov/about-data/publications/panimmune). MDSC infiltration, tumor dysfunction, and tumor exclusion scores were downloaded from the TIDE: Tumor Immune Dysfunction and Exclusion database (tide.dfci.harvard.edu/download). Genomic data and DNA repair metrics are available from Knijenburg et al. (Supplementary file “TCGA_DDR_Data_Resources.xlsx”).

## Supplementary Figures

**Figure_S1:**
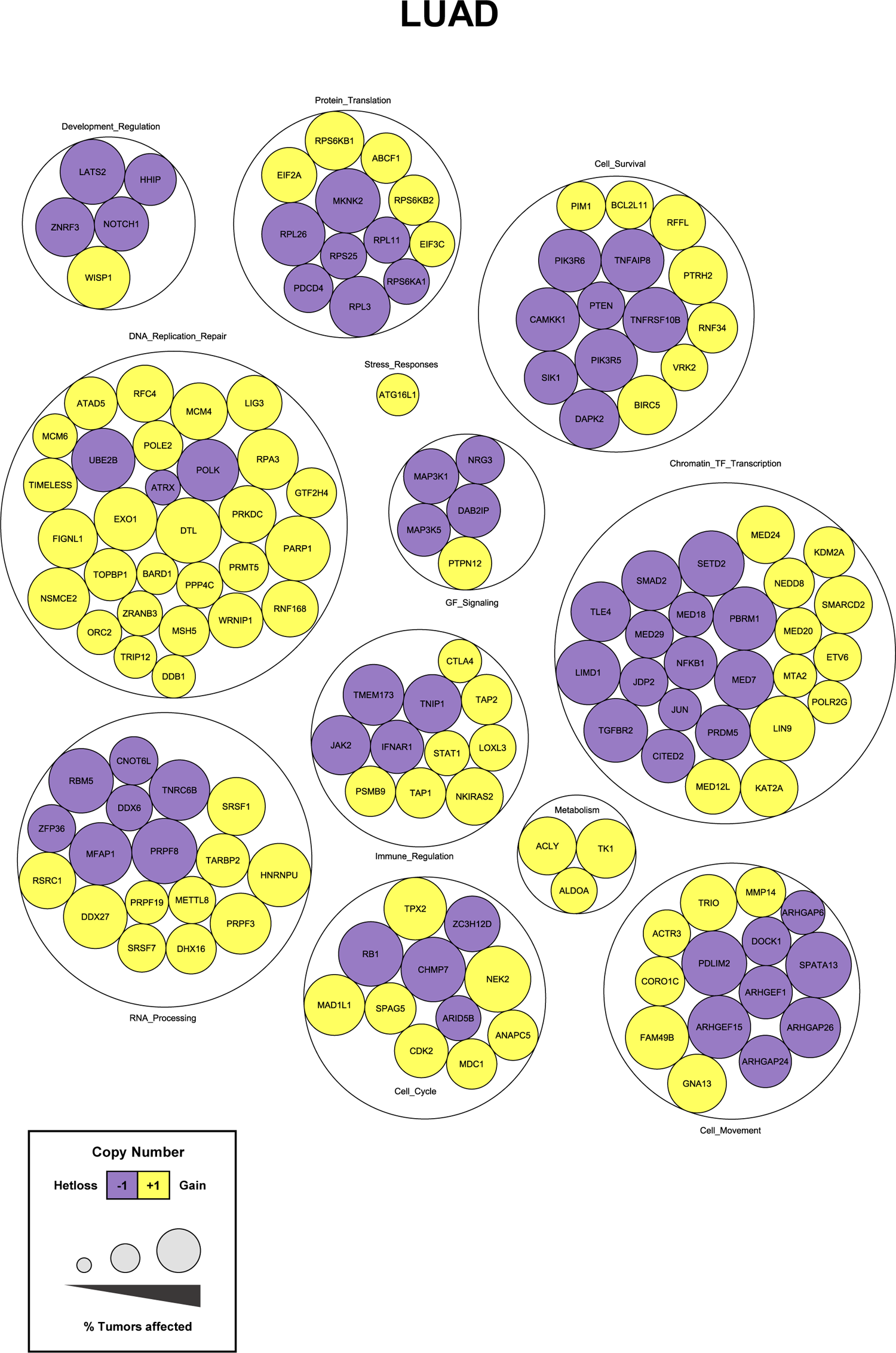
MYBL2 High lung adenocarcinoma replication stress sensitive site labeled functional cluster analysis.

**Figure_S2:**
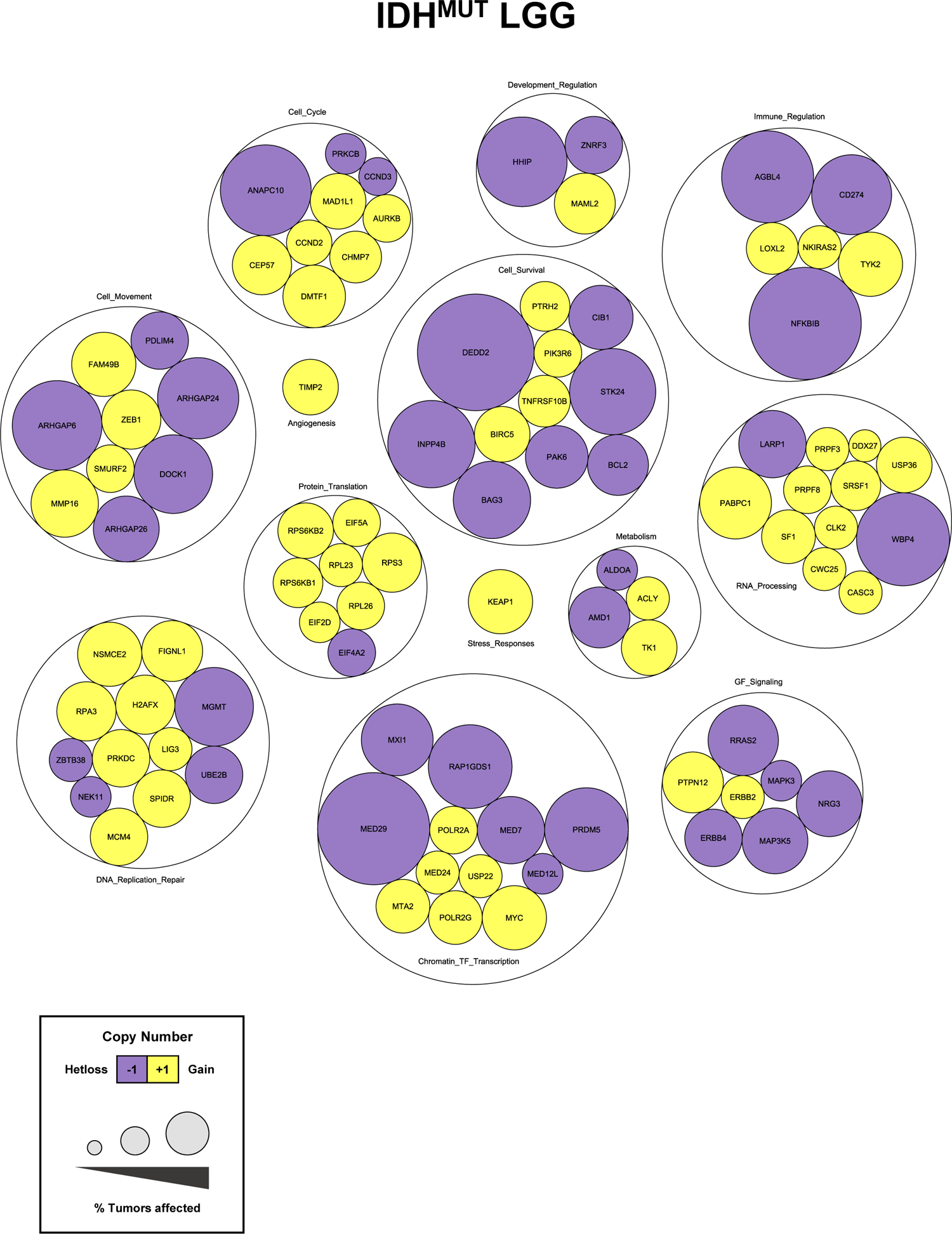
MYBL2 High IDH-mutant lower grade glioma replication stress sensitive site labeled functional cluster analysis.

**Figure_S3:**
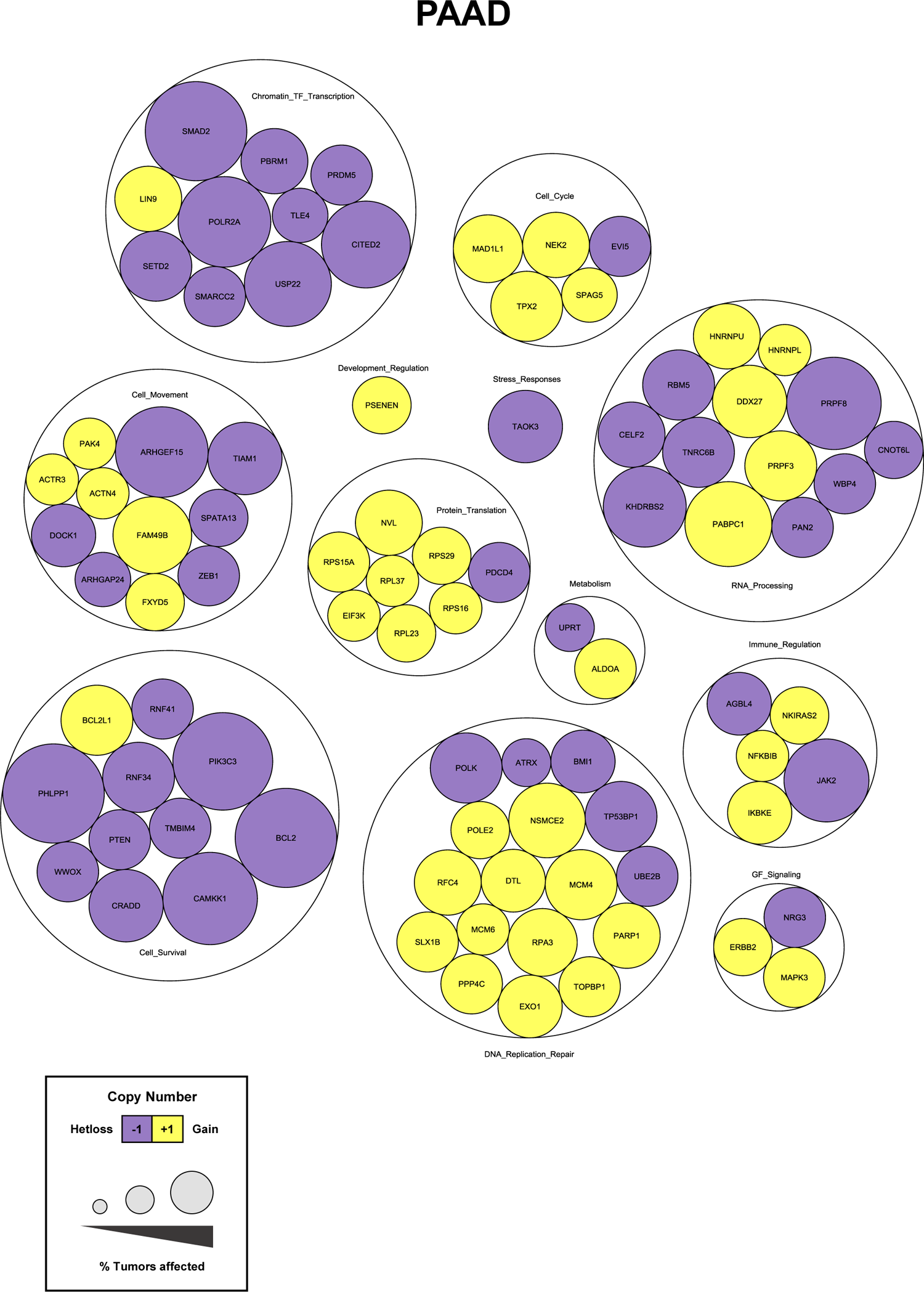
MYBL2 High pancreatic adenocarcinoma replication stress sensitive site labeled functional cluster analysis.

**Figure_S4:**
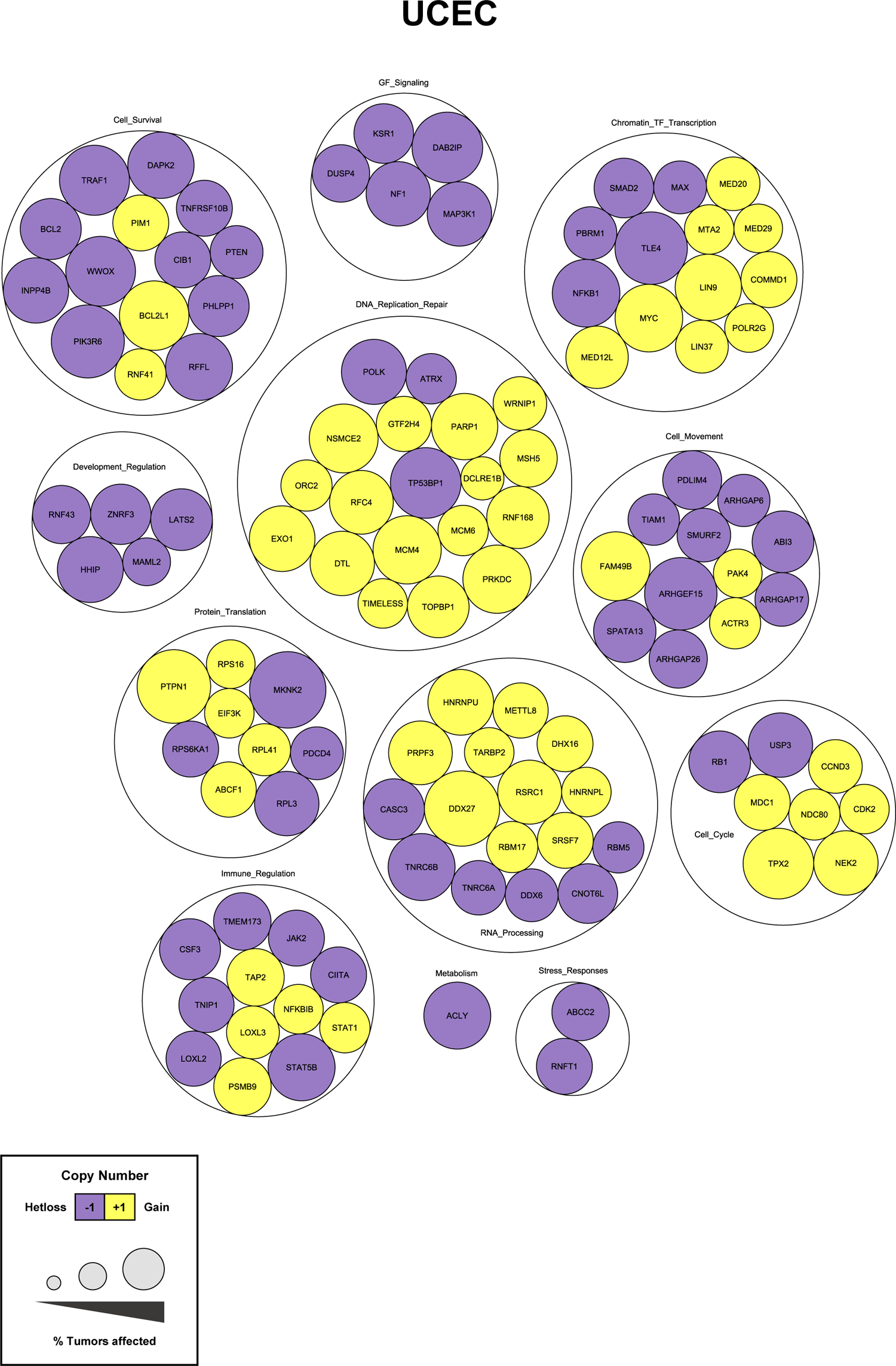
MYBL2 High endometrial carcinoma replication stress sensitive site labeled functional cluster analysis.

**Figure_S5:**
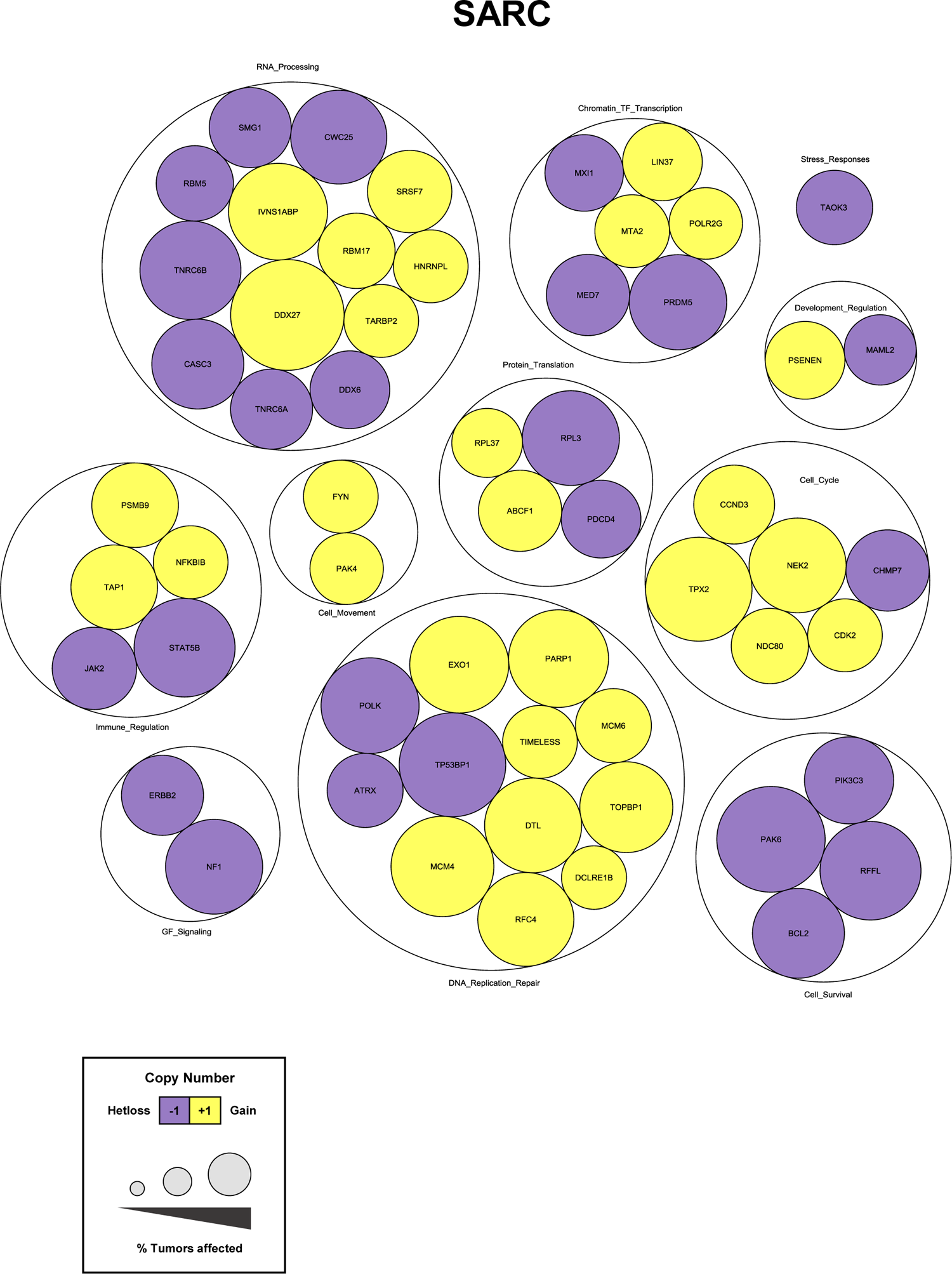
MYBL2 High sarcoma replication stress sensitive site labeled functional cluster analysis.

**Figure_S6:**
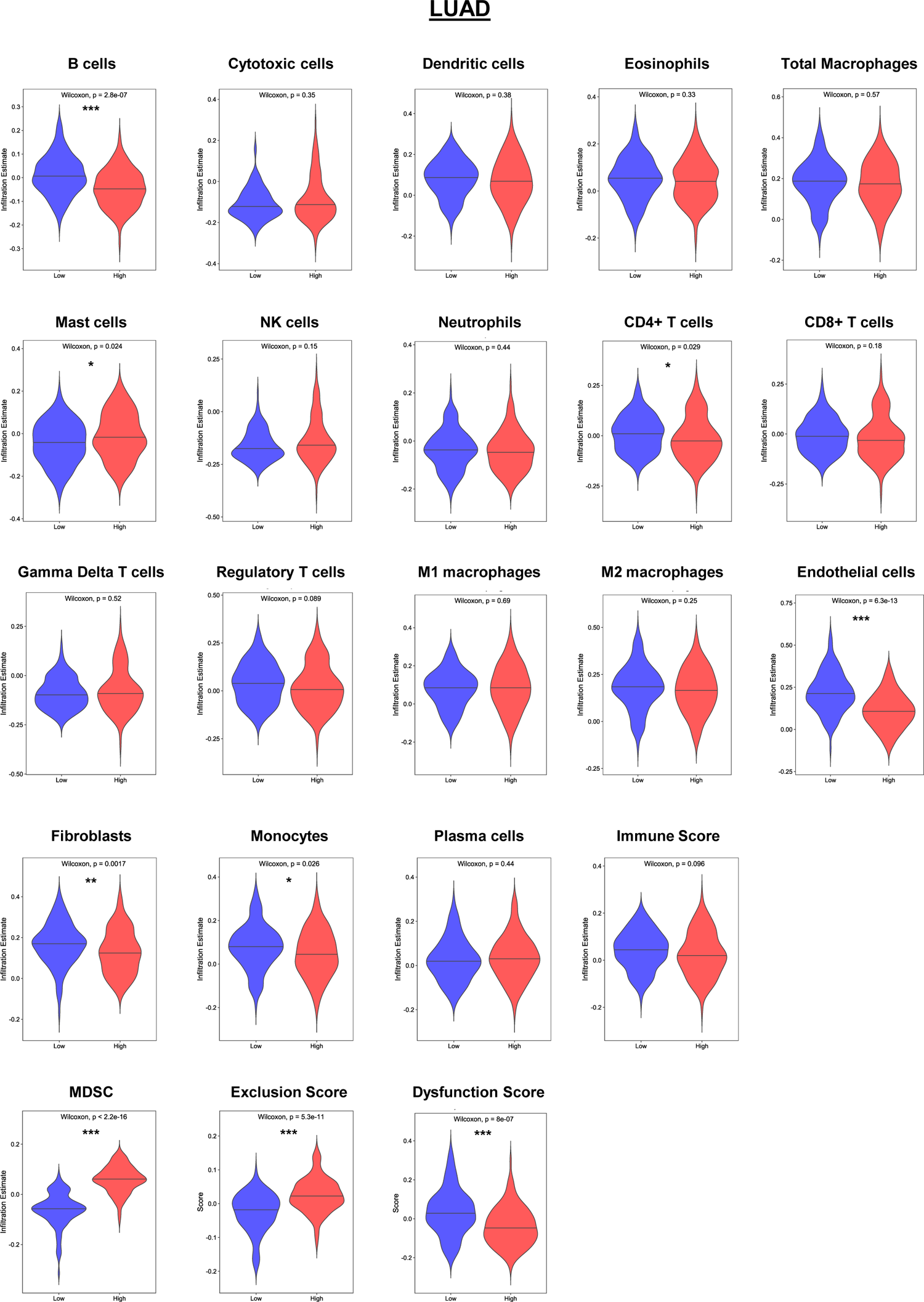
Lung adenocarcinoma ConsensusTME and TIDE analysis.

**Figure_S7:**
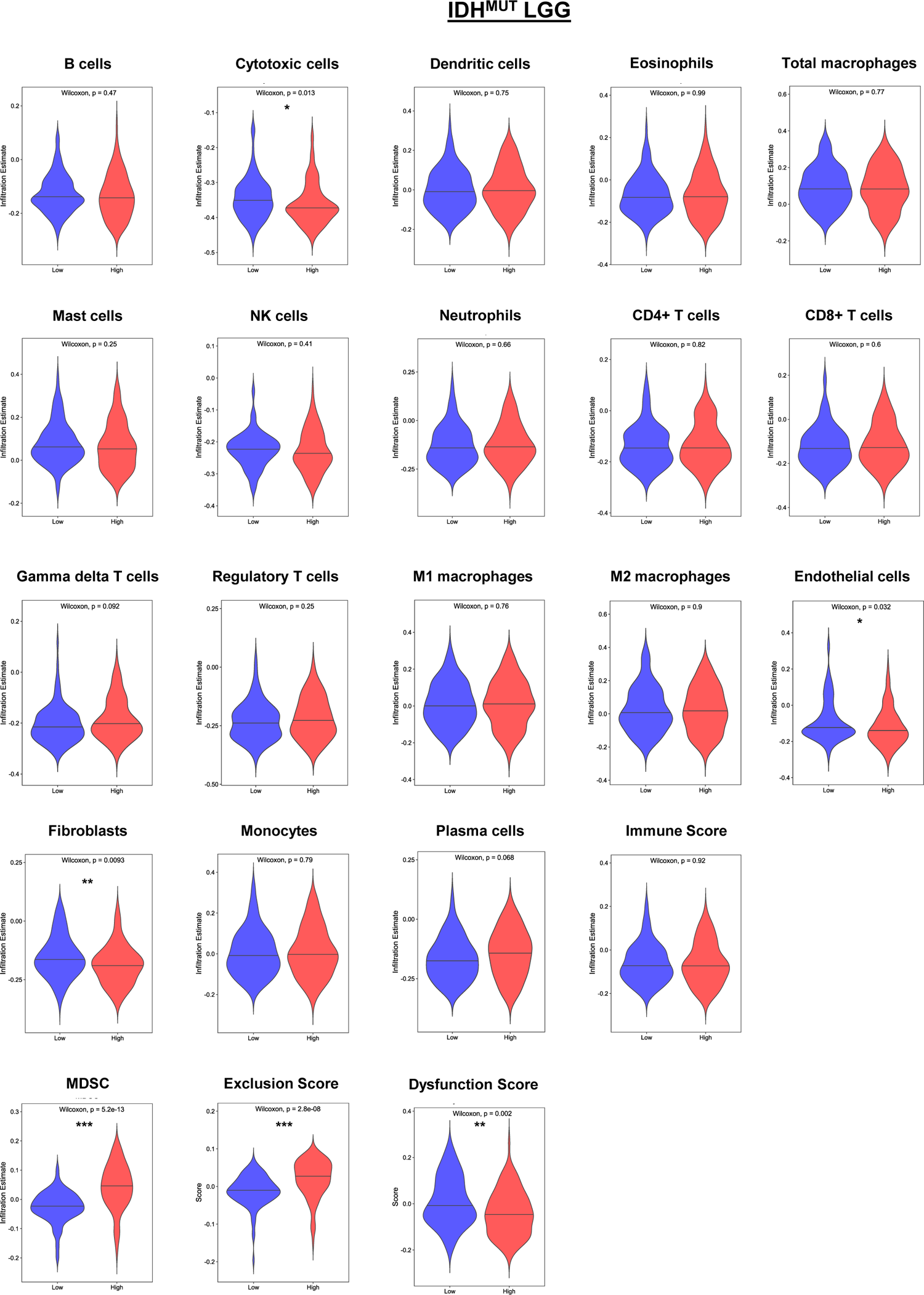
IDH-mutant lower grade glioma ConsensusTME and TIDE analysis.

**Figure_S8:**
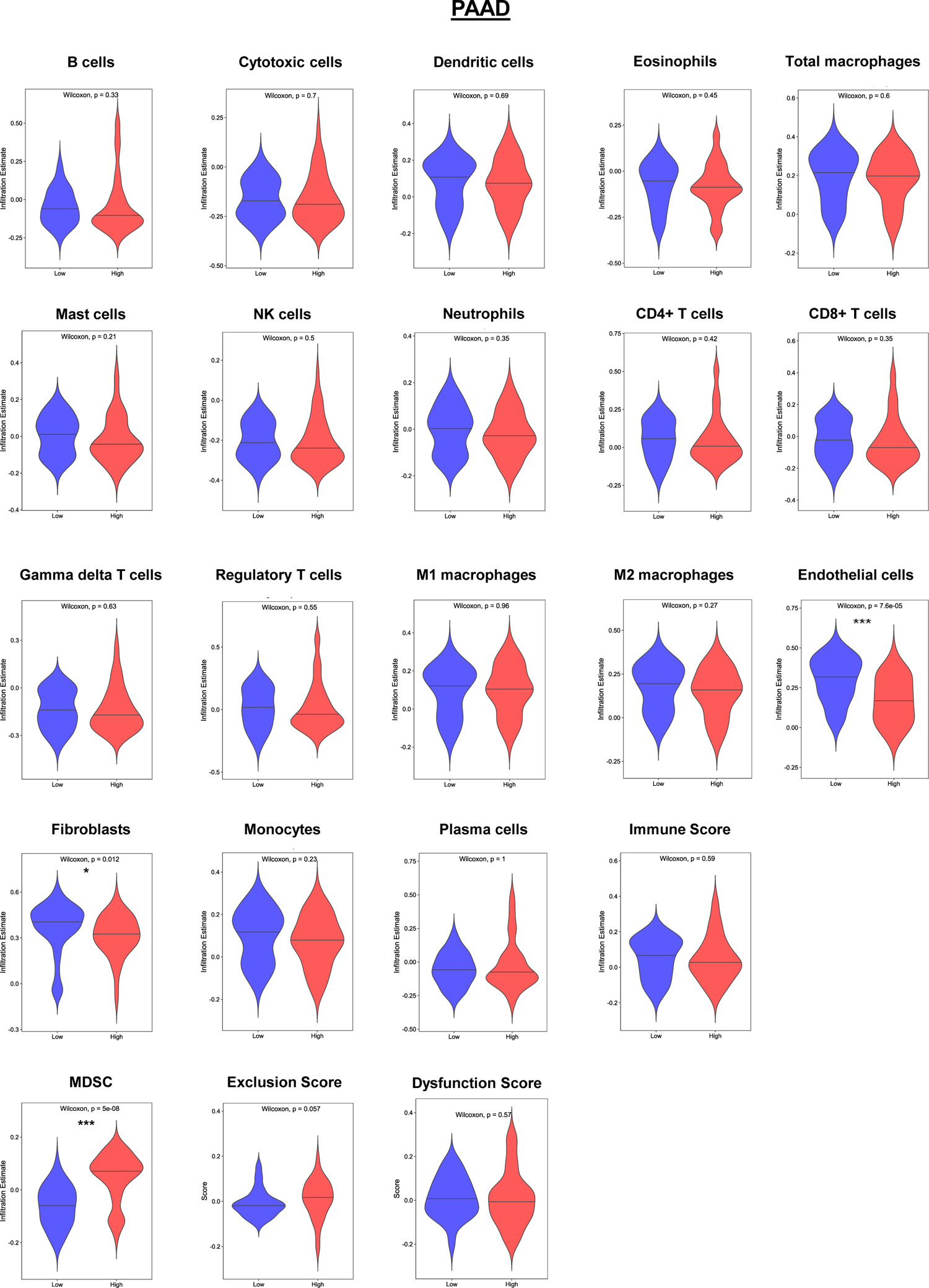
Pancreatic adenocarcinoma ConsensusTME and TIDE analysis.

**Figure_S9:**
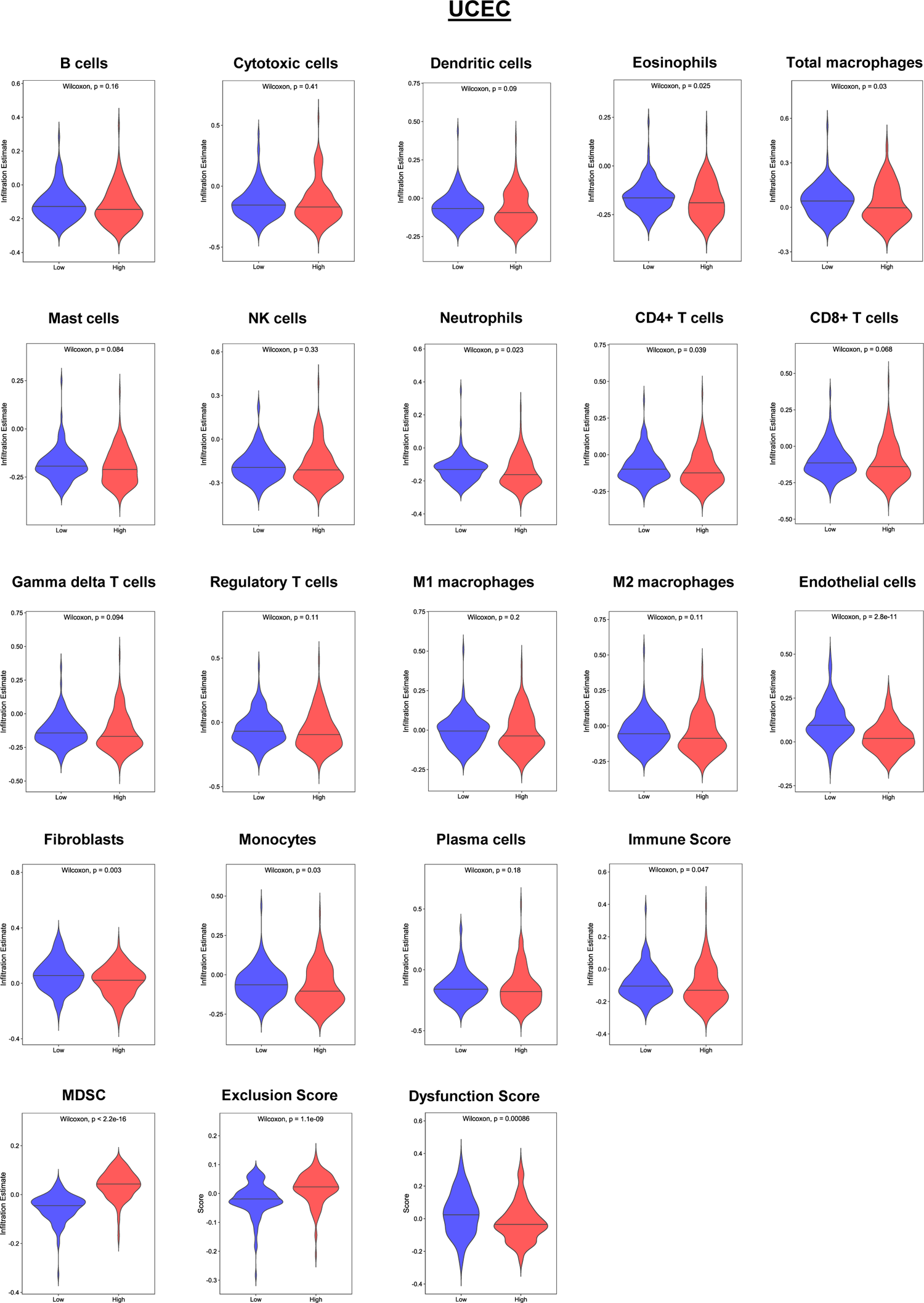
Endometrial carcinoma ConsensusTME and TIDE analysis.

**Figure_S10:**
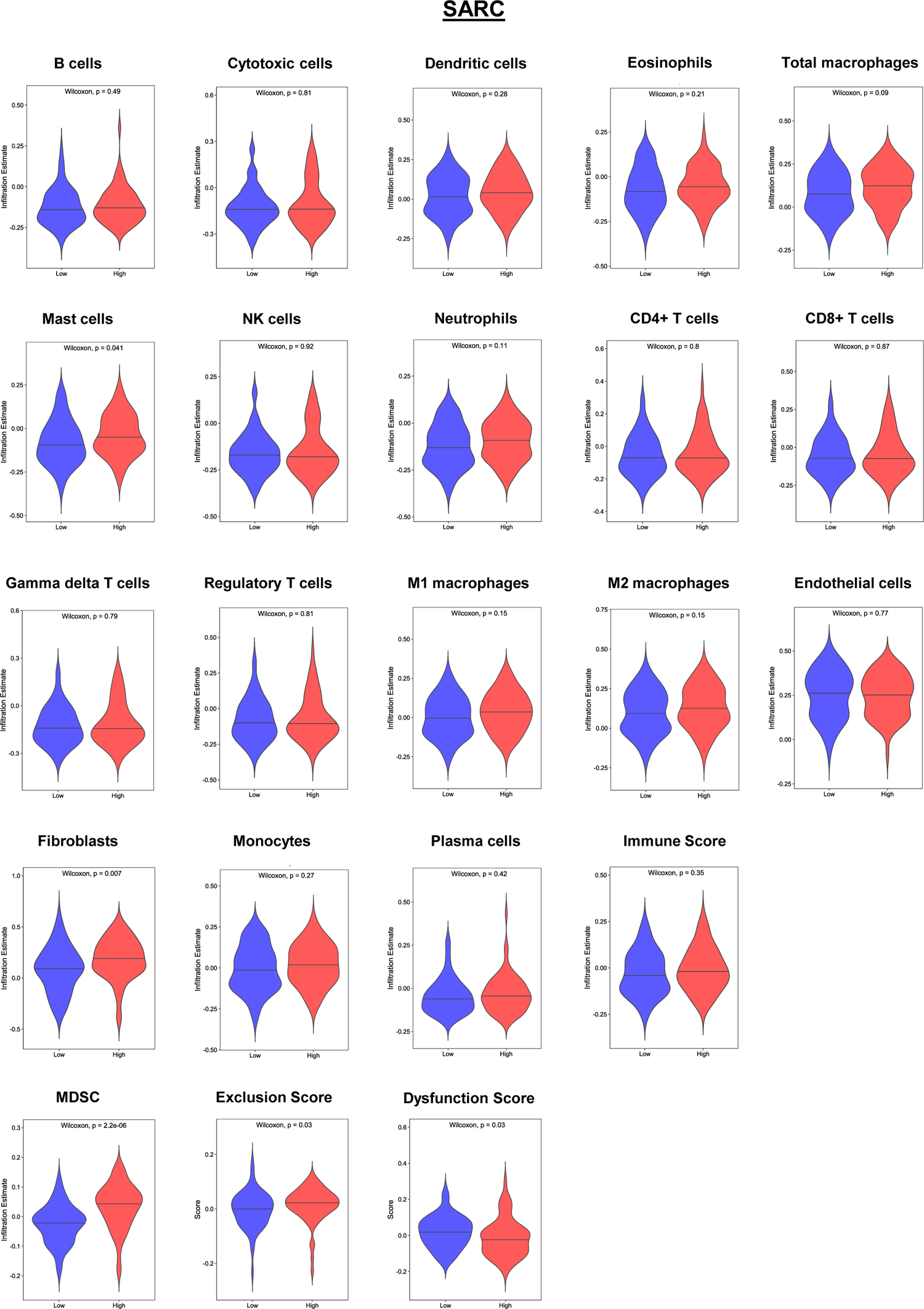
Sarcoma ConsensusTME and TIDE analysis.

**Figure_S11:**
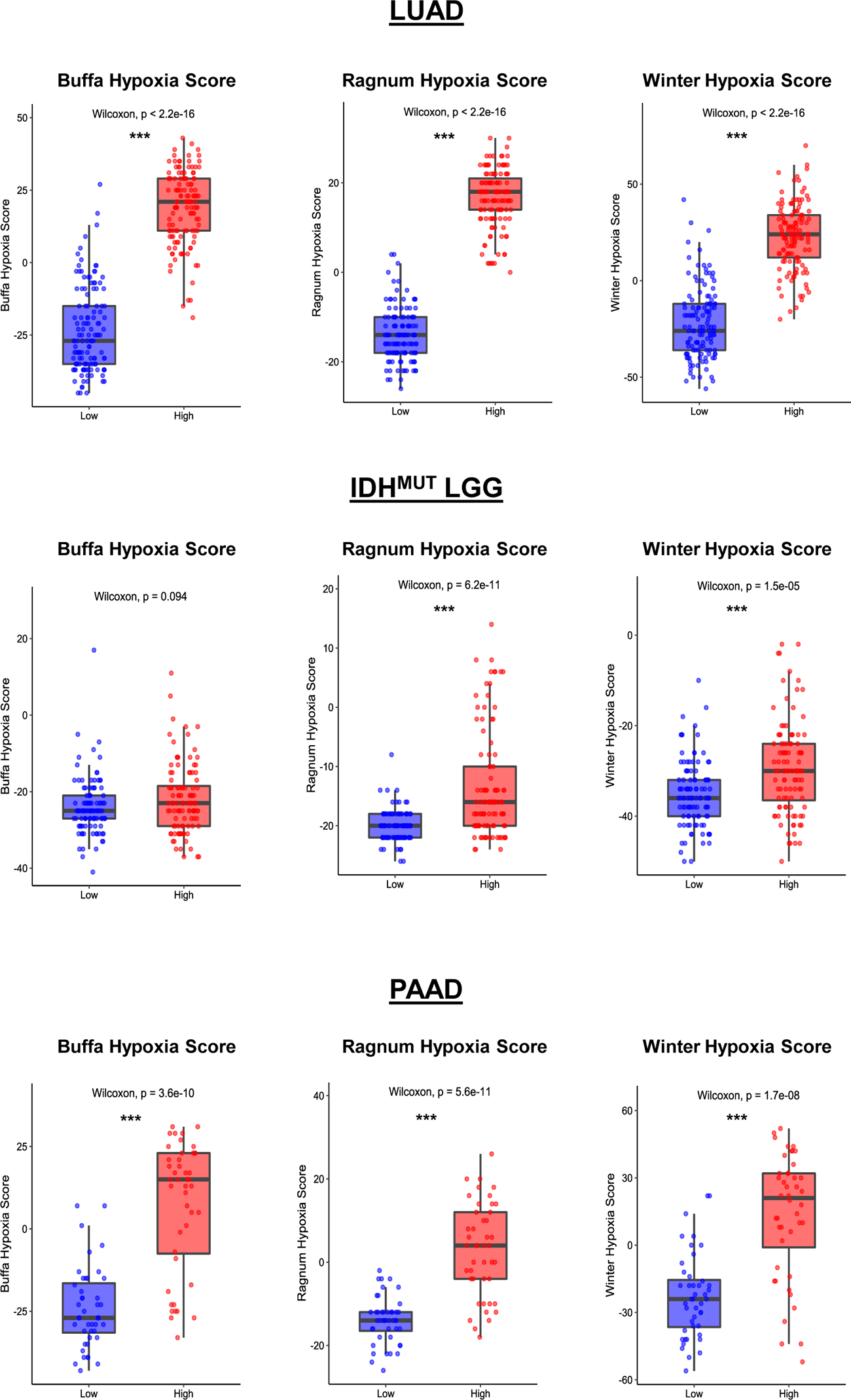
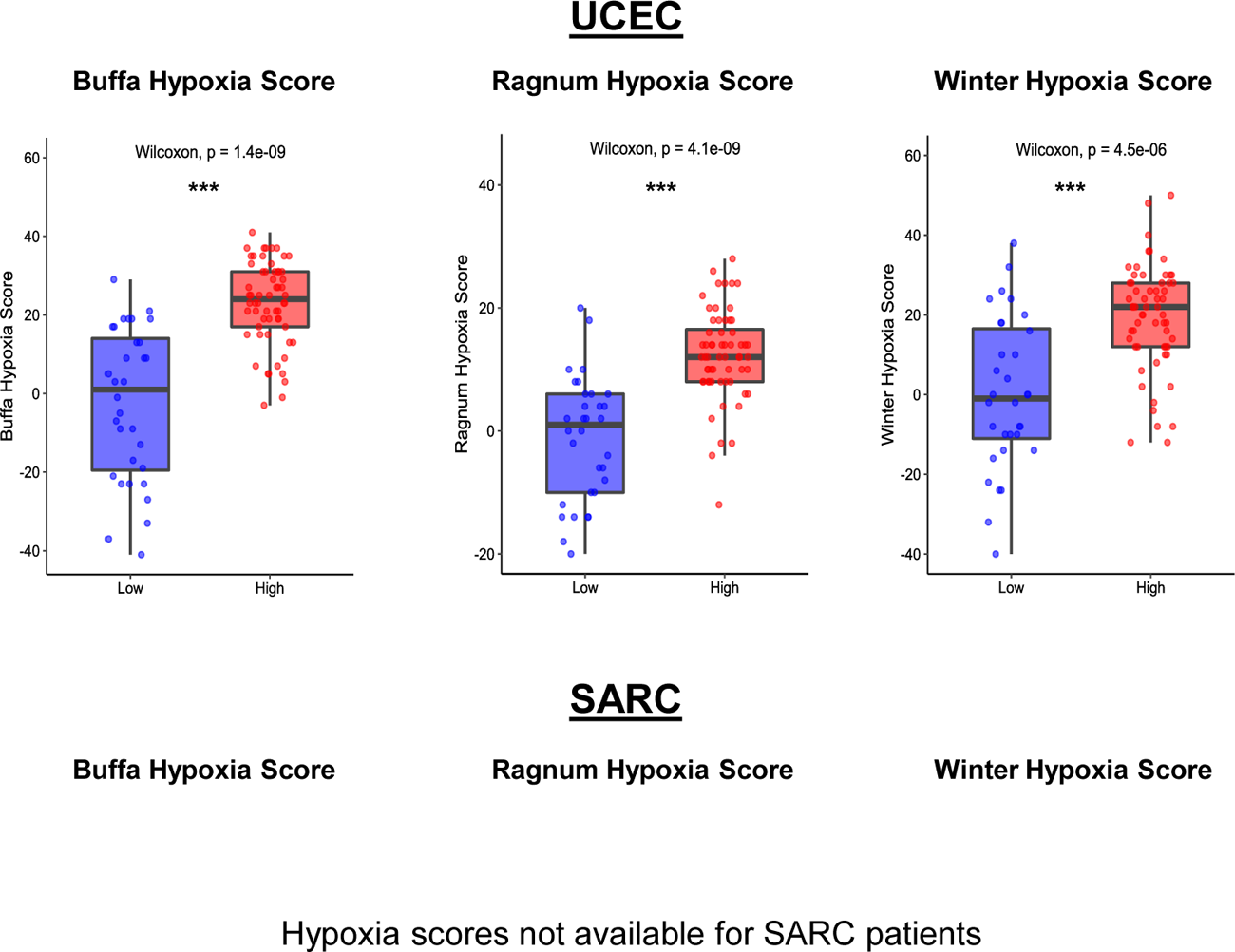
Hypoxia score analysis.

**Figure_S12:**
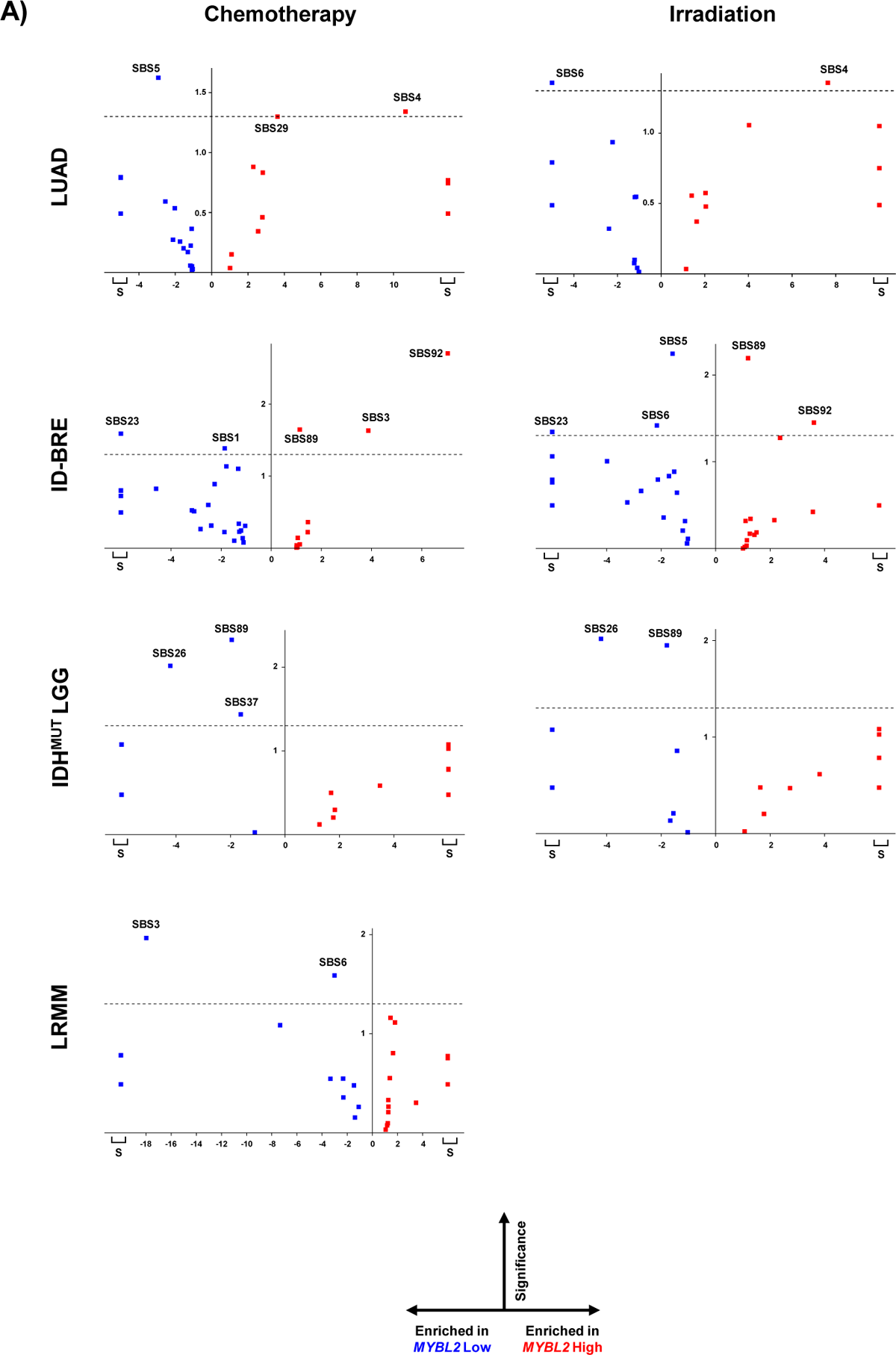
ORIEN COSMIC SBS 3.2 analysis.

**Figure_S13:**
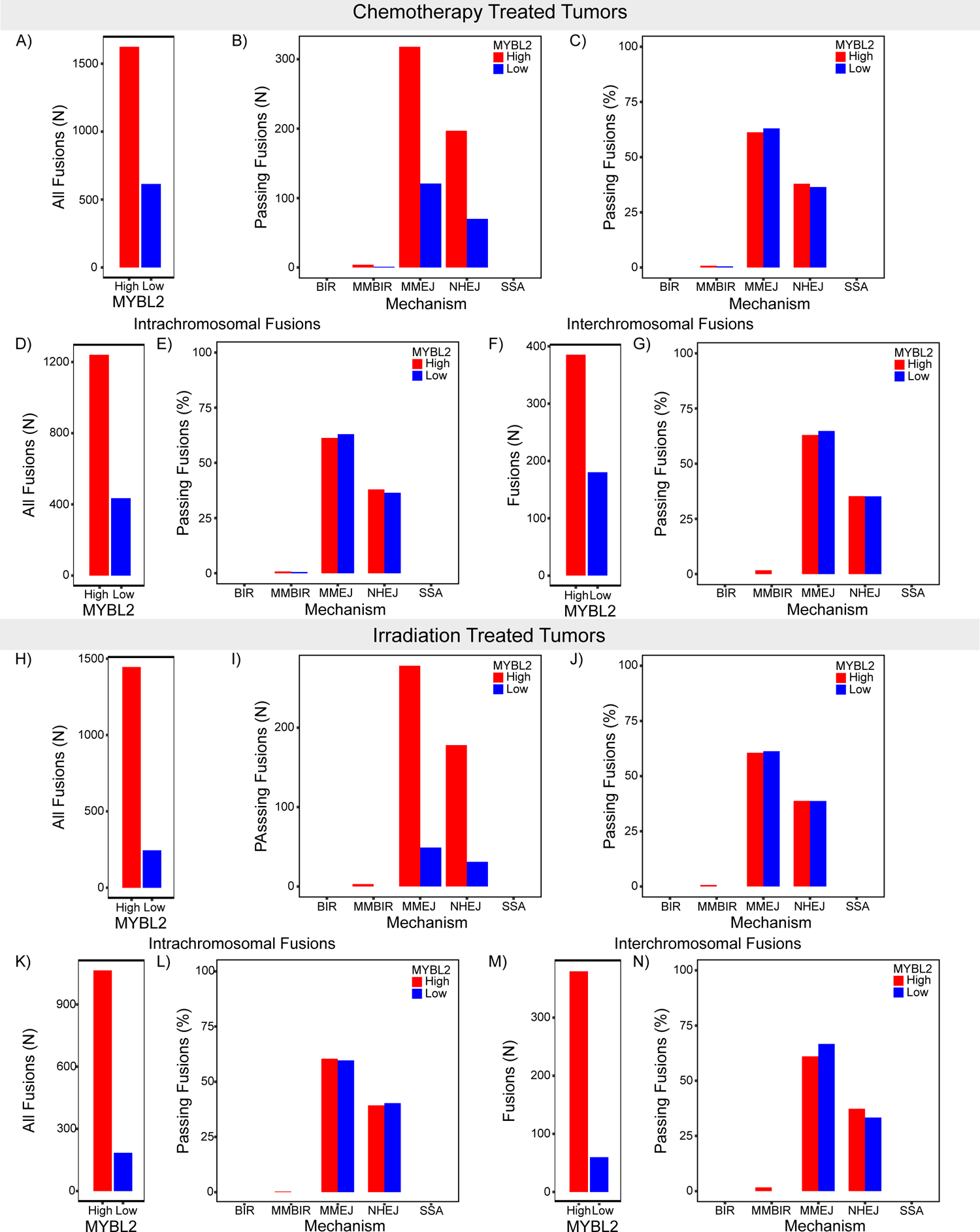
ORIEN LUAD *MYBL2* High Low therapy FUSED analysis.

**Figure_S14:**
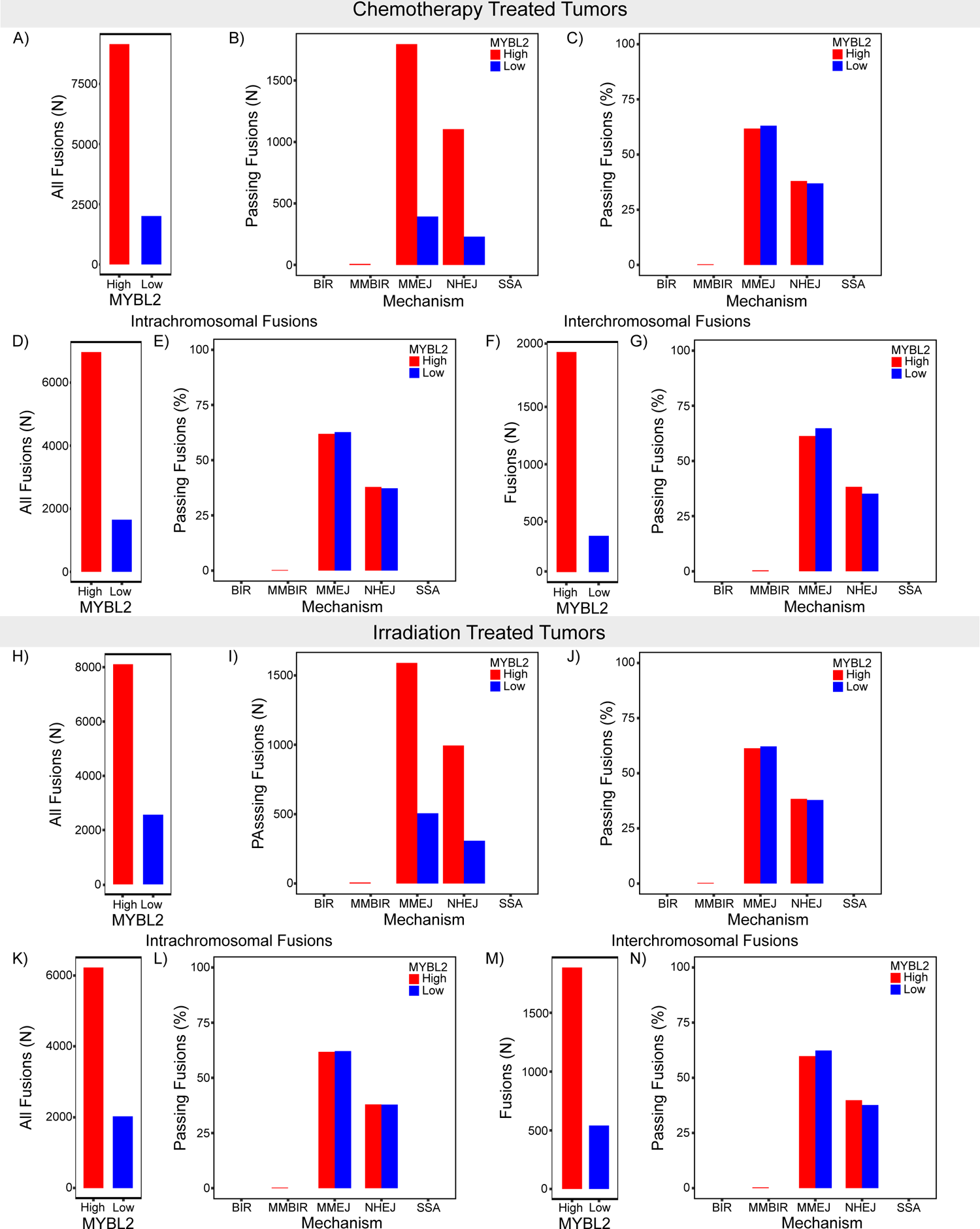
ORIEN ID-BRE *MYBL2* High Low therapy FUSED analysis.

**Figure_S15:**
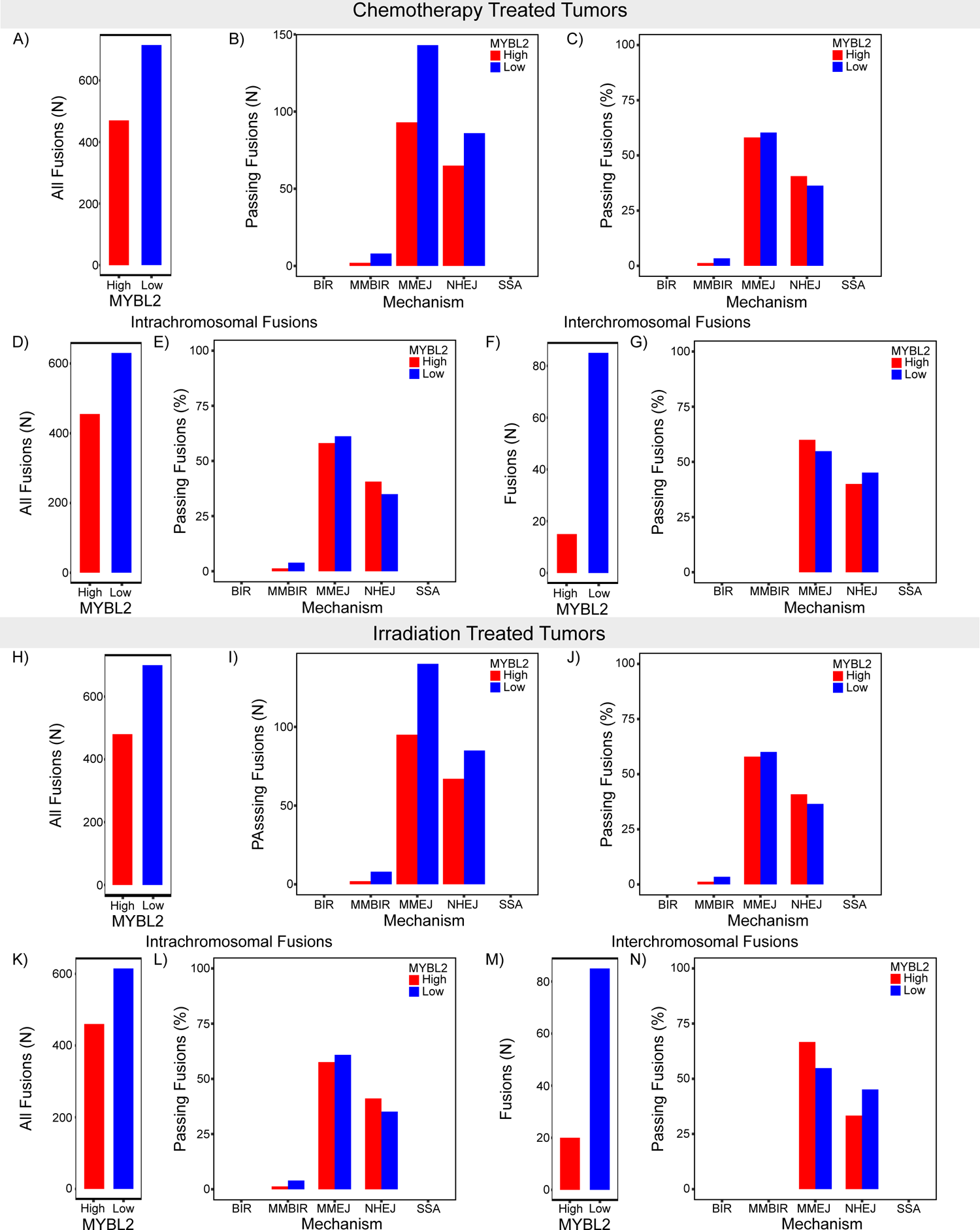
ORIEN IDH^MUT^ LGG *MYBL2* High Low therapy FUSED analysis.

**Figure_S16:**
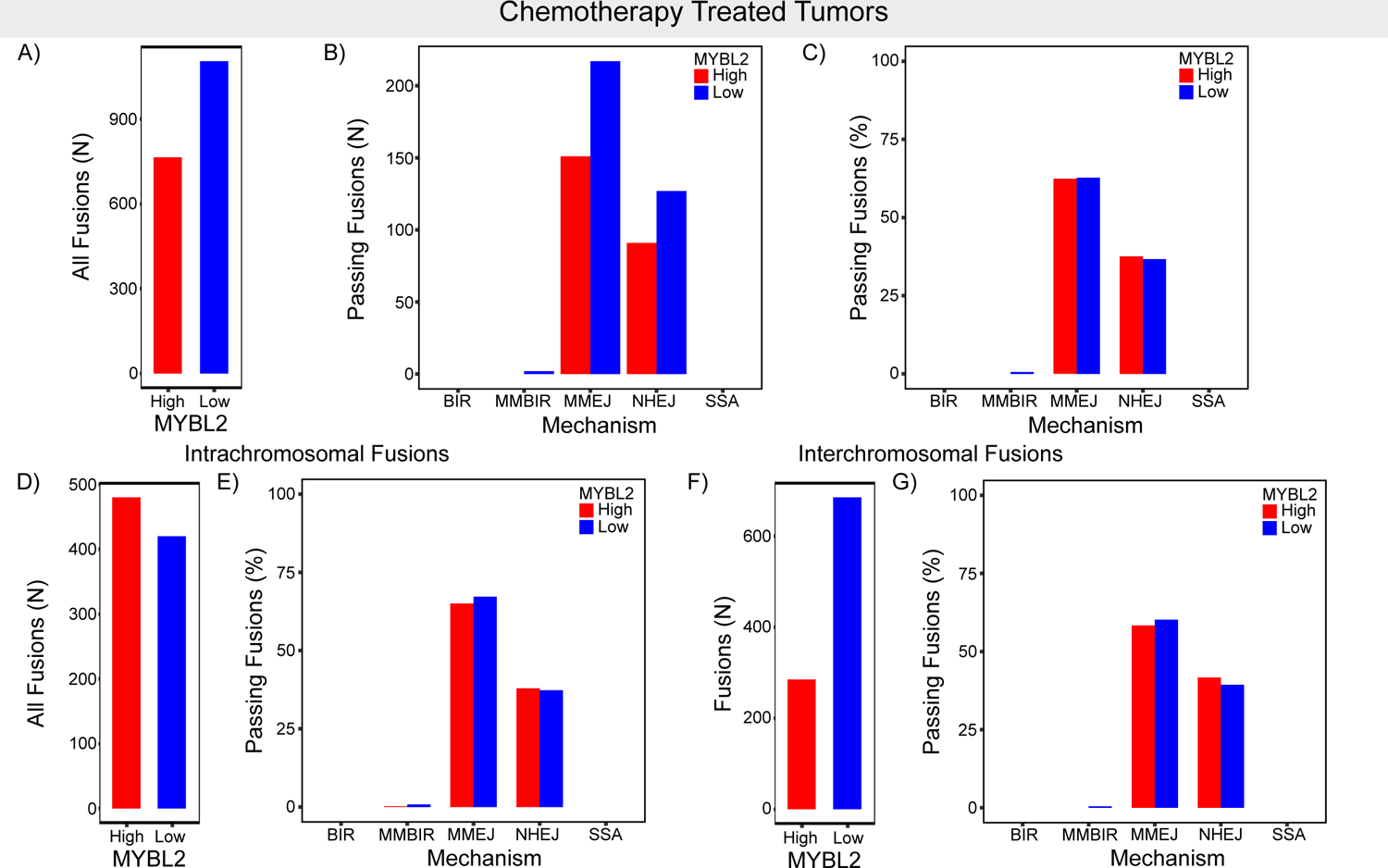
ORIEN LRMM *MYBL2* High Low therapy FUSED analysis.

**Figure_S17:**
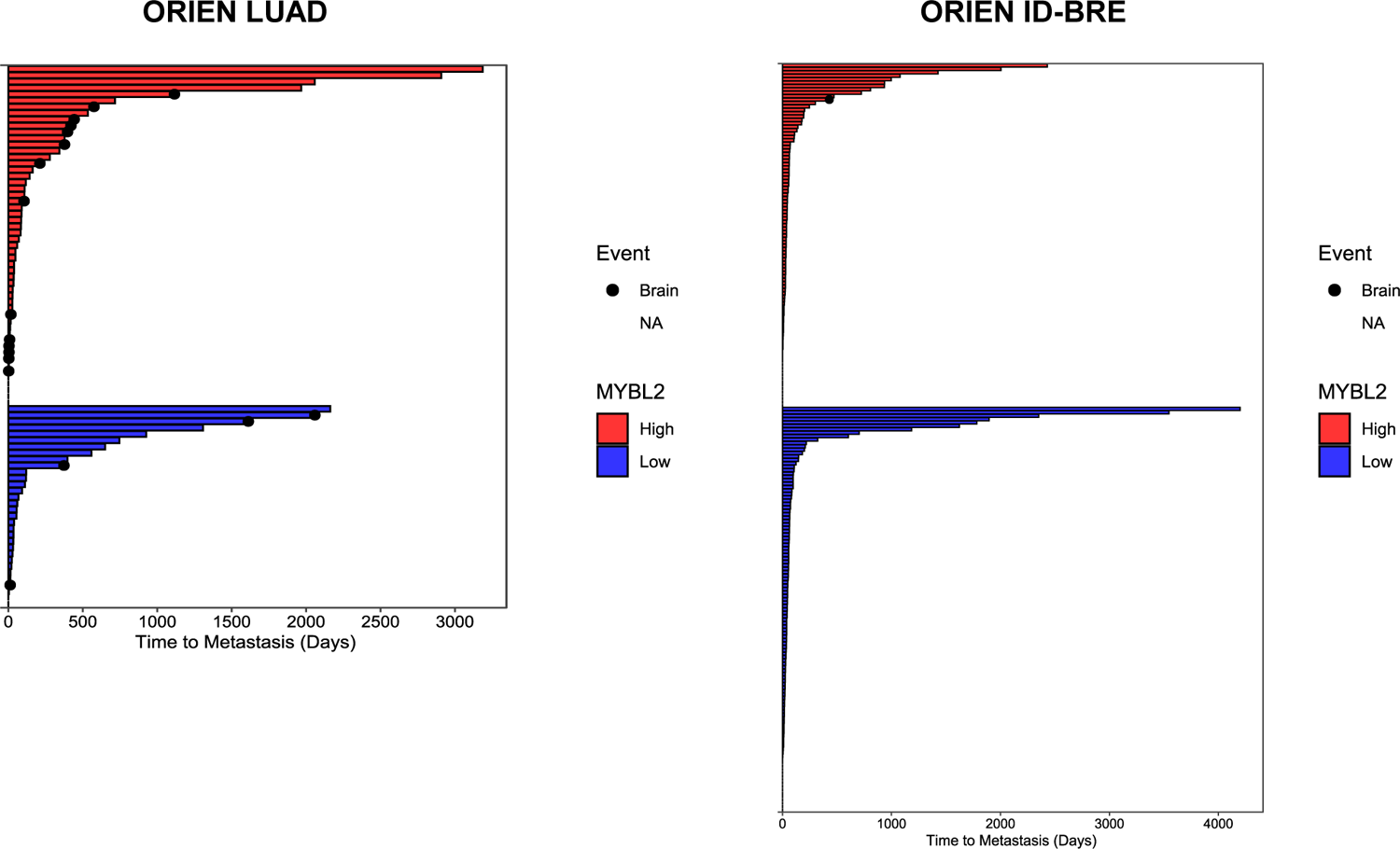
ORIEN time to metastasis swimmer plots.

## Supplementary Data

**Supplementary_Table_1**: Excel sheet detailing log-rank *p* values for *MYBL2* High vs. *MYBL2* Low OS, DSS, PFS outcomes across tumor types.

**Supplementary_Table_2**: Excel sheet containing final *MYBL2* High and Low patient identifiers and clinical data for all five tumor types.

**Supplementary_Table_3**: Excel sheet containing WE pathway score data for *MYBL2* High vs *MYBL2* Low tumors.

**Supplementary_Table_4:** Excel sheet containing replication stress score genes.

**Supplementary_Table_5**: Excel sheet containing replication stress sensitive site function cluster annotation.

**Supplementary_Table_6**: Excel sheet containing final replication stress sensitive site copy number alteration percentages along with RNA-seq differential expression values for all tumor cohorts.

**Supplementary_Table_7**: Excel sheet containing patient cohort numbers for TCGA and ORIEN Kaplan-Meier survival analyses in Figures 1 and 7

